# Regional variation in cholinergic terminal activity determines the non-uniform occurrence of cortical slow-wave activity during REM sleep

**DOI:** 10.1101/2022.03.01.481863

**Authors:** Mojtaba Nazari, Javad Karimi Abadchi, Milad Naghizadeh, Edgar J. Bermudez-Contreras, Bruce L. McNaughton, Masami Tatsuno, Majid H. Mohajerani

**Affiliations:** Canadian Centre for Behavioral Neuroscience, University of Lethbridge, Lethbridge, AB T1K 3M4, Canada; Center for Neurobiology of Learning and Memory, Department of Neurobiology and Behavior, University of California, Irvine, CA 92697, USA

**Keywords:** Sleep, non-rapid eye movement (NREM), rapid eye movement (REM), slow waves, hippocampus, neocortex, Cholinergic projection, acetylcholine, wide-field mesoscale voltage and glutamate imaging

## Abstract

Sleep consists of two basic stages: non-rapid eye movement (NREM) and rapid eye movement (REM) sleep. NREM sleep is characterized by slow high-amplitude cortical EEG signals, while REM sleep is characterized desynchronized cortical rhythms. While, until recently, it has been widely believed that cortical activity during REM sleep is globally desynchronized, recent electrophysiological studies showed slow waves (SW) in some cortical areas during REM sleep. Electrophysiological techniques, however, have been unable to resolve the regional structure of these activities, due to relatively sparse sampling. We mapped functional gradients in cortical activity during REM sleep using mesoscale imaging in mice, and observed local SW patterns occurring mainly in somatomotor and auditory cortical regions, with minimum presence within the default mode network. The role of the cholinergic system in local desynchronization during REM sleep was also explored by calcium imaging of cholinergic terminal activity within the mouse cortex. Terminal activity was weaker in regions exhibiting SW activity more frequently during REM sleep. We also analyzed Allen Mouse Brain Connectivity dataset and found that these regions have weaker cholinergic projections from the basal forebrain.

## Introduction

The brain has two well-characterized general behavioral states, waking and sleeping, with the sleeping state further divided into two phases: rapid eye movement (REM) sleep and non-REM (NREM) sleep. During the waking state, the brain receives sensory stimuli from the external world, processes it in the context of previous experience, and makes decisions and executes actions, if necessary. During the sleeping state, the brain is largely disconnected from the external world, but current understanding is that sleep is not just a passive state. Rather, it is an active state that is involved in several important functions, possibly including synaptic hemostasis and consolidation of memory^1,2^.

Both wakefulness and REM sleep are characterized by ‘desynchronized’, small-amplitude cortical EEG activity, and, at least in rodents, strong rhythmic activity in the hippocampus (theta waves). In contrast, during NREM sleep the cortex exhibits ‘synchronized’, large-amplitude slow-wave activity (SWA), and the hippocampus exhibits irregular bursts of high frequency spiking activity known as sharp-wave-ripples (SWR). These patterns of activity can also be observed under certain anesthetic agents such as urethane^3,4^. It has been widely accepted that these cortical activity patterns occur globally; however, a growing body of recent evidence has suggested that they can also occur locally^5–9^. During a waking state after a period of sleep deprivation, local cortical slow waves were found in awake humans^7^ and rats^6,8^. During NREM sleep, depth EEG recordings in humans^5^ and rodents^6^ have shown that slow-wave activity tends to recruit a limited number of cortical regions and form travelling waves that may not necessarily propagate across all the cortex^10,11^.

Recently, laminar electrophysiological recording in naturally sleeping mice suggested the presence of synchronized cortical activity in REM sleep, which was confined to local cortical regions and layers^9^. Although this finding provided an important insight into cortical dynamics during REM sleep, it was not possible to obtain a clear picture of their regional variation due to the sparse spatial sampling of the electrophysiological technique. Another recent study using EEG recording demonstrated the presence of local SWA in the primary sensory and motor cortices of humans during REM sleep^12^. However, because of volume conduction effects, the SWA could not be definitively localized to specific cortical regions. To overcome these limitations, we performed wide-field optical imaging of the whole mouse cortex during natural sleep, as well as under urethane anesthesia, which creates a brain state similar to natural sleep^3,4^.

Local synchronized cortical patterns occurred both during REM sleep and the urethane induced REM-like state, and they were more likely to occur in somatomotor and auditory cortical regions. Interestingly, these cortical networks negatively correlate with the Default Mode Network^13^ that is involved in the exchange of information between the hippocampus and the neocortex during memory recall^14,15^. We also investigated the role of the cholinergic system in regional cortical dynamics during REM sleep, using both structural and functional imaging of cholinergic axon and terminal activity within the whole cortex. Our results illustrate that regions with more SWA during REM sleep have weaker cholinergic projections from basal forebrain and lower levels of acetylcholine release. Furthermore, we demonstrated that increasing the level of extracellular acetylcholine, significantly reduces the occurrence of slow wave in REM sleep. These findings shed light on possible mechanisms underlying local modulation of cortical activity and their role in information processing during sleep.

## Results

### Glutamate imaging of brain activity during sleep in head-restrained mouse

Glutamate imaging of spontaneous cortical activity was performed in Ai85-CamKII-Emx mice, which express the glutamate sensing fluorescent reporter iGluSnFR in excitatory cortical neurons^16^, while the mice slept under head-fixation (Fig. 1a, supplementary Fig. 1a,b). Glutamate imaging provides sufficient spatiotemporal resolution (50 um, 100 Hz) for capturing cortical SWA with a large field of view, and it also allows for chronic skull-intact imaging without performing a craniotomy^14^. The primary challenge of this type of recording, however, is that mice do not tend to fall asleep while head-restrained. To overcome this difficulty, we maintained the temperature of both the room and the recording platform at a higher level, and minimized sounds and vibrations, in order to allow the animals to fall asleep in the recording setup. Animals were habituated to the recording setup for at least 2 weeks prior to imaging. To increase the success rate of sleep recording, animals were gently sleep restricted the day before the recording (see methods). Glutamate imaging was combined with electrophysiological recording of hippocampal CA1 local field potentials (LFP) and neck muscle activity (supplementary Fig. 1c). These electrophysiological signals were used to score sleep stages and wakefulness (methods). To further assist with scoring, pupil diameter of the animals was also measured (Fig. 1). Spontaneous activity was recorded in several ~30-minute sessions, which was sufficient to capture wake to sleep transitions (~58% of recording sessions had sleep events). The implanted unilateral imaging window was large enough to include multiple cortical regions (Fig. 1b). Active waking was characterized by high muscle tone, hippocampal theta activity, and large pupil size^14,17^(Fig. 1c).

**Figure 1.**
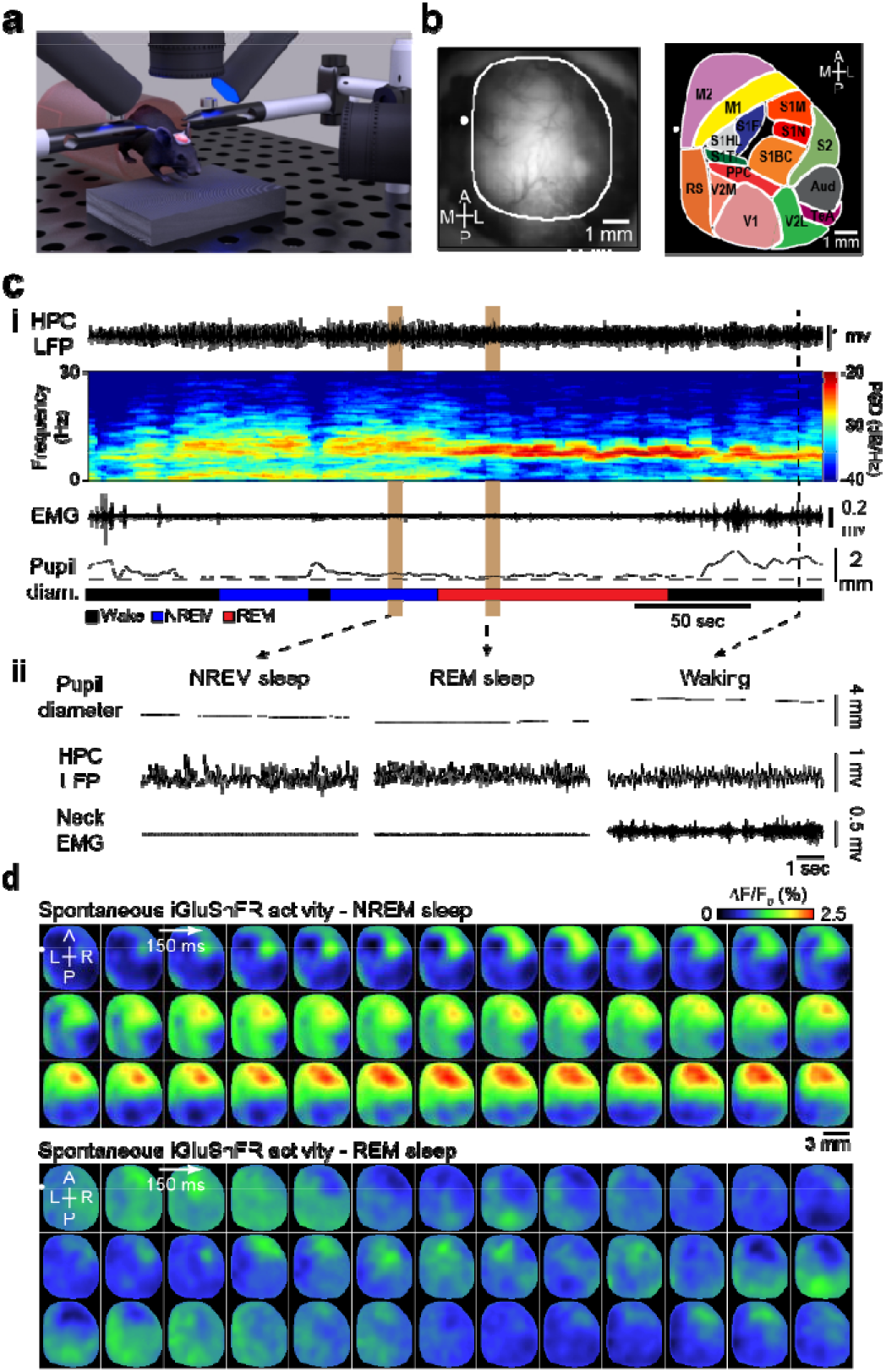
Longitudinal glutamate imaging of brain activity during sleep in head restrained animal. (a) Experimental setup for mesoscale glutamate imaging of head-fixed mouse. (b) Cranial window preparation of widefield glutamate imaging (left) and a cartoon representing imaged cortical regions (right). (c) (i) Traces of hippocampal LFP, neck muscle EMG and pupil diameter during states of wakefulness, NREM sleep and REM sleep in a representative head-fixed mouse. These traces were used to score sleep and waking states. Power spectrogram of the hippocampal LFP shows clear transitions across slow wave activity in NREM sleep, theta activity in REM sleep and wakefulness. (ii) Close-up of LFP, EMG and pupil signals in three behavioural states. (d) Montage of spontaneous glutamate activity corresponding to two epochs of NREM and REM sleep states highlighted in c.

NREM sleep was characterized by low electromyography (EMG) activity, high-amplitude irregular hippocampal activity, and oftentimes small pupil size. REM sleep was characterized by low muscle tone, rhythmic hippocampal theta activity, and very small pupil size (Fig. 1c, supplementary Fig. 2a,b). The peak frequency and bandwidth of REM sleep theta activity were similar between head-fixed sleep-restricted mice and unrestrained mice (see methods)(supplementary Fig. 2c). However, the former group showed significantly longer REM sleep episodes and inter-REM intervals (supplementary Fig. 2d, p-value < 0.01, Kolmogorov-Smirnov test). This is consistent with studies showing increased sleep pressure after sleep restriction^18,19^.

Transitions between the three behavioural states was reflected in the spectral profile of the iGluSnFR signal. For example, slow (< 1 Hz) and delta band (1-4 Hz) power for two cortical regions (secondary motor cortex (M2) and primary visual cortex (V1)) were reduced in REM sleep compared to NREM sleep (supplementary Fig. 2e). This was in line with the fact that the neural population activity, modulating the glutamate fluorescence signal, transitions from irregular phasic firing during NREM sleep to more regular, tonic firing during REM sleep^20,21^. An exemplar montage of images illustrates the cortex-wide propagation of depolarization waves during the up/down oscillations in NREM sleep. In REM sleep, the reduction of the cortical glutamate signal is apparent; however, some localized large-amplitude activities are still present (Fig. 1d).

### Local cortical synchronization during REM sleep

We assessed the spatial extent of local slow-wave activity that occurred in REM sleep. During REM sleep, the cortex exhibited desynchronized, small-amplitude activity throughout the cortical mantle; however, sometimes, certain regions exhibited synchronized activity (Fig. 2a). To assess the spatial profile of cortical synchronization in REM sleep, we calculated the combined slow-wave and delta power during REM and NREM sleep at pixel level. P-value maps were constructed at each pixel by statistically comparing NREM and REM sleep power distributions using the Wilcoxon rank-sum test (see methods). Based on the p-value map, the cortex was often divided into two intracortical network: one that transitioned to a significantly desynchronized state during REM sleep (p < 0.05, Fig. 2b right panel, dark colour), and the other that continued to exhibit synchronized activity during REM sleep (p ≥ 0.05, Fig. 2b right panel, hot colour).

**Figure 2.**
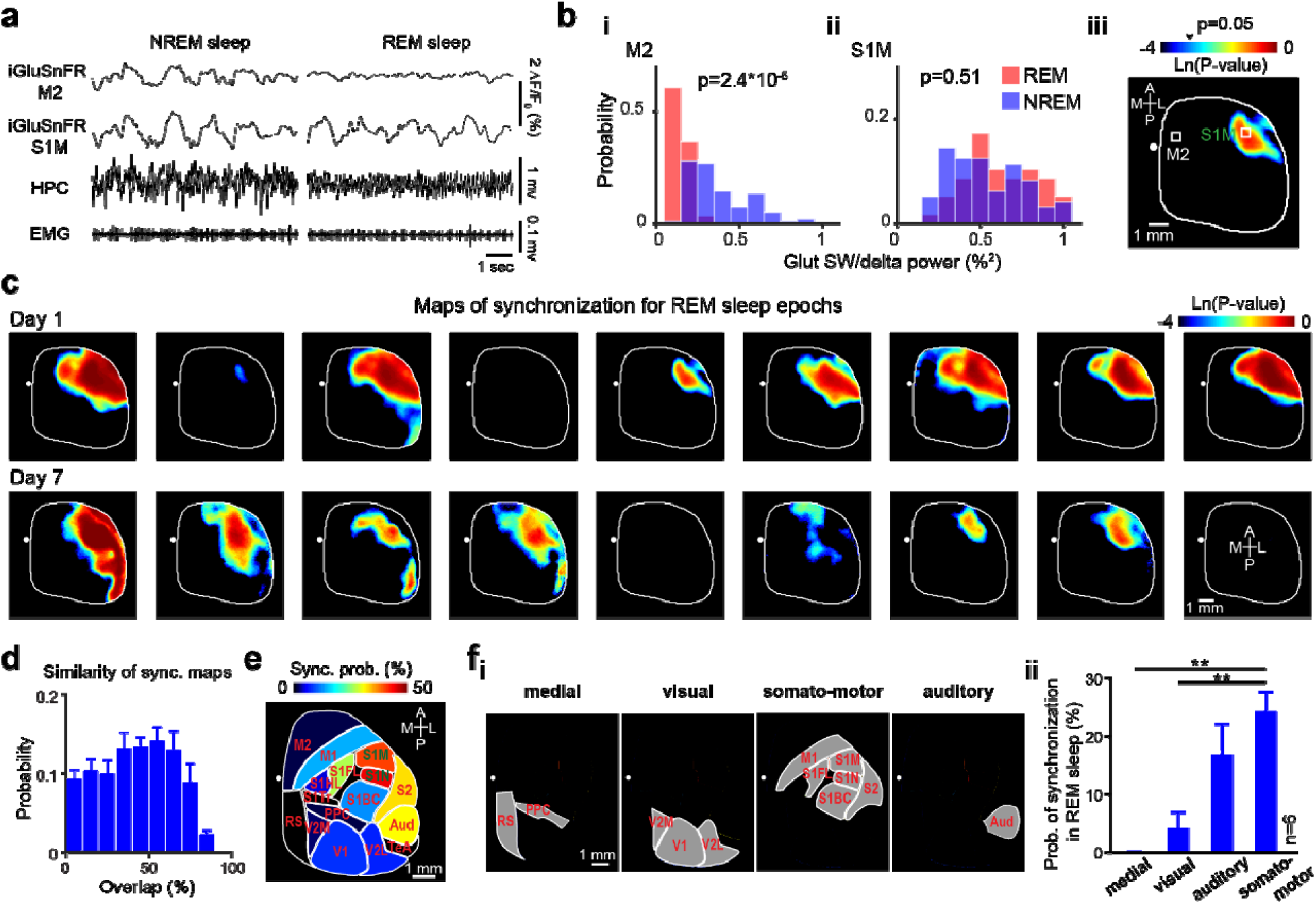
Local synchronization of cortical activity in REM sleep. (a) An example of iGluSnFR filtered signal (0.2-5 Hz) derived from ROIs (0.112 mm^2^) in secondary motor cortex (M2) and mouth primary somatosensory (S1M), as well as hippocampal LFP and EMG signals during an epoch of NREM (left) and REM (right) sleep. Note that S1M area shows slow-wave activity in both NREM and REM sleep. (b) Histograms showing distribution of iGluSnFR signal power (0.5-5 Hz) during an epoch of REM (red) and NREM (blue) sleep in M2 (i) and S1M (ii). P-values are calculated from comparing NREM and REM sleep power distributions using Wilcoxon rank-sum test. (iii) P-value map is calculated from comparing power distributions for each pixel within the entire imaging field. The pixel intensities are scaled logarithmically. (c) Representative p-value maps calculated for 9 non-overlapping epochs (8 sec long) of a REM sleep episode across two different days. Note that patterns of p-value maps are heterogeneous and change across cortex over different REM epochs. (d) Histogram of overlap between all pairs of binarized p-value maps pooled across 6 mice. Error bars, s.e.m. (e) Binarized p-value maps were averaged and registered onto the Allen Institute Mouse Brain Coordinate Atlas (n=6 animals). Warmer color means higher probability of synchronization or presence of slow-wave activity during REM sleep. (f) (i) Four major structurally defined cortical subnetworks. (ii) Bar graph shows probability of synchronization during REM sleep across cortical subnetworks, sorted in ascending order. Error bars, s.e.m. (*P < 0.05, **P < 0.01, Repeated measure ANOVA with Greenhouse-Geisser correction for sphericity: F3,15 = 17.947, p = 0.0024; post-hoc multiple comparison with Tuckey’s correction: medial vs visual p = 0.5869, medial vs auditory p = 0.1146, medial vs somatomotor p = 0.0050, visual vs auditory p = 0.0487, visual vs somatomotor p = 0.0026, auditory vs somatomotor p = 0.1449).

Interestingly, the cortical regions exhibiting synchronized activity appears to be localized within anterior-lateral sections of imaging field of view, they were not uniformly distributed across the cortex over different REM sleep epochs (Fig. 2c). We quantified this observation by calculating the overlap of the spatial profile of cortical synchronization between the p-value maps (see methods); if the spatial profiles of cortical synchronization are stable, the overlap should be skewed to 100%. Instead, the overlap values were skewed to 0% (p-value < 0.001, Lilliefors test), with a median of 39.3%, indicating that the spatial profile of cortical synchronization are relatively heterogeneous over different REM sleep epochs (Fig. 2d). To quantitatively measure the probability of synchronization during REM sleep, we averaged binarized p-value maps and registered them onto the Allen Mouse Brain reference atlas, based on anatomical landmarks (bregma and lambda) and sensory-evoked mapping (see methods) (Fig. 2e). Out of all the imaged regions, mouth and nose primary somatosensory cortices (S1M, S1N) showed the highest probability of synchronization during REM sleep (Fig. 2e and Supplementary Fig. 3a). While there is much variability in the initiation regions from mouse to mouse (Fig. 2d), regions exhibiting synchronized activity appear to be located on the midline, as well as distributed both medially and posteriorly. Surprisingly, these areas overlap with cortical regions that are activated around the hippocampal sharp-wave ripple (SWR) complex (Supplementary Fig. 3b)^14^. Cortical area activated by SWR is centered around the Default Mode Network as defined in the mouse brain^13,22^.

Interestingly, regions with comparable synchronization probability tended to fall within previously identified cortical subnetworks^23^. The somatomotor subnetwork that includes forelimb, mouth, and nose primary somatosensory, secondary somatosensory, and primary motor cortices along with the auditory subnetwork stayed synchronized during REM sleep for the longest, followed by visual and medial networks (Fig. 2g; n = 6 per subnetwork; repeated measure ANOVA with Greenhouse-Geisser correction for sphericity: F3,15 = 17.947, p = 0.0024; post-hoc multiple comparison with Tuckey’s correction: medial vs visual p = 0.5869, medial vs auditory p = 0.1146, medial vs somatomotor p = 0.0050, visual vs auditory p = 0.0487, visual vs somatomotor p = 0.0026, auditory vs somatomotor p = 0.1449). In addition to calculating the p-value maps, we also calculated the SW/delta activity power across different regions during REM sleep. We confirmed that somatomotor and auditory regions also show the strongest SW/delta power compared to other regions during REM sleep (Supplementary Fig. 3b,c).

Since previous work^24^ has suggested that local SWA and phasic motor activity during REM are correlated, we contrasted the presence of local SWA between phasic and tonic REM sleep. To do so, phasic REM events were separated from tonic ones (Supplementary Fig. 4a). The face movement signal was used to separate phasic REM events from tonic ones. The p-value maps for phasic and tonic REM epochs were calculated in the six Ai85-CamKII-EMX mice strain used for glutamate imaging. In contrast to previous studies^24^, we found that the probability of SWA presence were highly correlated between phasic and tonic REM sleep across 14 cortical regions (r =0.96, paired t-test, p>0.05 for all comparisons, Supplementary Fig. 4b,c).

### Electrophysiological recording in unrestrained sleeping mice showed local SWA during REM sleep

In addition to glutamate imaging of head-restrained mice, we investigated the local occurrence of synchronization using electrophysiological recording from unrestrained sleeping mice. Bipolar electrodes were implanted in primary and secondary motor, retrosplenial, barrel, and mouth primary somatosensory cortices of eight C57 mice (Fig. 3a). Some of these cortical areas showed synchronized, and some showed desynchronized activity during REM sleep in glutamate imaging animals. Hippocampal and EMG electrodes were also implanted for scoring the behavioural state. LFP recording from these mice started one hour after the onset of the light cycle (8:30 AM) and continued for 4 hours. This was repeated for 3 days. Wakefulness and NREM/REM sleep were scored based on hippocampal LFP and EMG.

**Figure 3.**
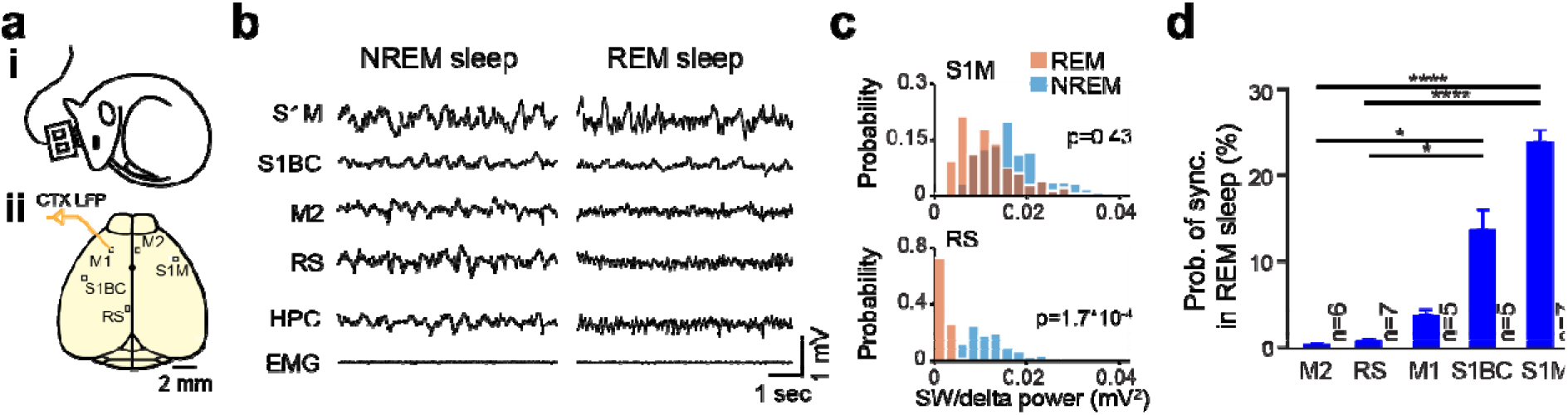
Multi-electrode LFP recording from unrestrained mouse during sleep shows that probability of synchronized activity during REM varies across cortical regions. (a) (i) Cartoon of sleep recording preparation in freely behaving mice. (ii) Schematic of the mouse cortex showing position of cortical electrodes from which the LFP signals were recorded. (b) Example of LFP and EMG signals during NREM and REM sleep. Note the local occurrence of slow-wave activity during REM sleep within S1M and S1BC. (c) Distribution of LFP SW/delta power (0.5-4 Hz) in NREM and REM sleep for two cortical regions: mouth primary somatosensory (S1M, top) and retrosplenial cortex (RS, bottom). Slow activity power was significantly reduced from NREM to REM sleep for RS while it is comparable for S1M (ranksum Wilcoxon test). (d) 5 recording sites are sorted by the probability of SWA presence during REM sleep. Error bars, s.e.m. (One-way ANOVA: F4,25 = 12.43, p = 1.059×10^−5^; post-hoc multiple comparison with Tuckey’s correction: secondary motor vs primary mouth p = 5.183×10^−5^, secondary motor vs primary barrel p = 0.0459, retrosplenial vs primary mouth p = 3.967×10^−5^, retrosplenial vs primary barrel p = 0.0474, primary motor vs primary mouth p = 7.988×10^−4^, p>0.1 for other comparison).

In agreement with the results of the glutamate recordings, SWA was present during REM sleep in some cortical regions’ LFPs (Fig. 3b). Following the p-value map calculation procedure, discussed before, we quantified the occurrence of synchronized activity in each cortical region by comparing the SW/delta power (0.5-4 Hz) of cortical LFPs between NREM sleep and REM sleep epochs (Fig. 3c). Similar to the glutamate imaging results, probability of synchronization was higher in primary mouth and barrel somatosensory cortices compared to the midline regions, including the secondary motor and retrosplenial cortices during REM sleep (Fig. 3d, One-way ANOVA: F4,25 = 12.43, p = 1.059×10^−5^; post-hoc multiple comparison with Tuckey’s correction: secondary motor vs primary mouth p = 5.183×10^−5^, secondary motor vs primary barrel p = 0.0459, retrosplenial vs primary mouth p = 3.967×10^−5^, retrosplenial vs primary barrel p = 0.0474, primary motor vs primary mouth p = 7.988×10^−4^, p>0.1 for other comparisons).

### Voltage imaging revealed local synchronization in urethane anesthetized mouse

Since iGluSnFR indicator measures concentration of extracellular glutamate^16^ which may not translate to membrane depolarization, for example due to dendritic inhibition, voltage sensitive dye (VSD) imaging was used to directly measure population membrane depolarization^25,26^. VSD signal has faster kinetics than iGluSnFR^27^, and its red spectral property mitigate neuroimaging artifacts due to flavoprotein autofluorescence, hemodynamics, respiration and heartbeat^28,29^. Mesoscale VSD imaging was performed under urethane anesthesia which causes the brain to cycle between NREM-like and REM-like states, mimicking the physiological correlates of the different stages observed during natural sleep^3,4^. Accordingly, seven C57 mice were anesthetized, implanted with hippocampal and cortical electrodes and imaged.

As reported before^3,4,30^, the NREM-like state was characterised by slow-frequency (~1 Hz) cortical LFPs (Supplementary Fig. 5a), whereas the REM-like state consisted of desynchronized cortical activity and hippocampal theta at a lower frequency than natural sleep (4.5±1.1 Hz, 7.4±0.3 Hz, and 7.5±0.2 Hz in urethane anesthesia (n=7 mice), head-restrained sleep (n=6 mice), and unrestrained sleep (n=8 mice) respectively; see Supplementary Fig. 2c for statistical comparisons). Hippocampal and cortical LFP state alternations coincided with a general reduction of VSD signal power during the REM-like state compared to the NREM-like state (supplementary Fig. 5a-d)^20,21^. Reduced power of the VSD signal in the REM-like state was observed in almost all of the frequency bands from 0.1 to 50 Hz, but most of the power reduction was concentrated in the frequencies less than 1 Hz (supplementary Fig. 5b). Reduction of the SW power can be seen as the absence of up/down state in the VSD montage when comparing NREM-like to REM-like states (supplementary Fig. 5f).

Cortical synchronization in the REM-like state was local, and these local patterns detected by the p-value maps changed between different REM-like epochs (supplementary Fig. 6a,b,c). Overlap of binarized p-value maps showed a non-normal distribution (p-value < 0.001, Lillieforc test), with a median of 37.5% (supplementary Fig. 6d), which is consistent with the glutamate recording (Fig. 2d). Similar to the glutamate sleep recording, probability of synchronization showed a heterogenous distribution across cortical regions (supplementary Fig. 6e). Group data analysis for all cortical regions indicates that somatomotor and auditory regions exhibited the highest probability of synchronized activity (supplementary Fig. 6f; n = 7 per subnetwork; repeated measure ANOVA with Greenhouse-Geisser correction for sphericity: F3,18 = 15.996, p = 0.0021; post-hoc multiple comparison with Tuckey’s correction: medial vs visual p = 0.0590, medial vs auditory p = 0.0379, medial vs somatomotor p = 3.779×10^−4^, visual vs auditory p = 0.0421, visual vs somatomotor p = 2.425×10^−3^, auditory vs somatomotor p = 0.9786).

These results also demonstrate that the probability of synchronized activity did not differ significantly between head-fixed REM sleep and the REM-like state observed under urethane anesthesia (t-test p-value>0.05, Supplementary Fig. 7a). In addition, across these cortical areas there was a significant positive correlation between the probability of synchronization in REM sleep and the REM-like state (Supplementary Fig. 7a; r=0.98, p<0.001). To directly compare glutamate sleep with urethane anesthetized results, three of the glutamate mice were recorded under urethane anesthesia after completion of the head-fixed sleep recording. Similar cortical maps of synchronization were found in both recording paradigms (Supplementary Fig. 7b, c and d; r=0.79, p<0.001).

### Investigation of acetylcholine’s role in non-uniform cortical desynchronization during REM sleep

It has been shown that the level of acetylcholine is high during REM sleep^31,32^ and the REM-like^4^ state and that elevated acetylcholine levels activate the neocortex, partly through disinhibition, and induce tonic firing of cortical neurons^33^. The basal forebrain (BF), the main source of cholinergic input to the cortex, is topographically organized based on its cortical projections^34,35^. There are also studies suggesting that the spatial distribution and laminar density of these projections differ across the cortex^36,37^. Accordingly, we hypothesized that the regions that receive weaker cholinergic innervation would show more SWA during REM sleep. To test this hypothesis, we assessed the regional distribution of cholinergic innervation received by cortex from the BF, using the Allen Mouse Brain Connectivity (AMBC) Atlas. This atlas provides projection mapping data that details the cortical cholinergic axonal innervation from different parts of the basal forebrain. These axonal projections were labeled with eGFP by injection of a Cre-dependent adeno associated viral (AAV) tracer into 10 BF nuclei sites in Chat-IRES-Cre mice^38^.

The AMBC mice had the same genetic background as the animals used for imaging and electrophysiology, and were age-matched. For each injection site, the 3D AMBC data matrix was converted to a 2D matrix by averaging the entries across the dimension associated with cortical layers (Fig. 4a, see methods for details). Next, all 2D maps were averaged across experiments to summarize the cholinergic innervation that the cortex receives from the BF (Fig. 4b). In the summary map, anterior-posterior axis of midline cortical regions showed dense cholinergic projections compared to lateral cortical regions. Finally, 14 cortical regions were registered on this map based on anatomical landmarks. The strength of BF cholinergic innervation in each region of interest (ROI) was negatively correlated with the probability of synchronized activity during the REM state (Fig. 4c; r=−0.79, p<0.001; n=10 experiments).

**Figure 4.**
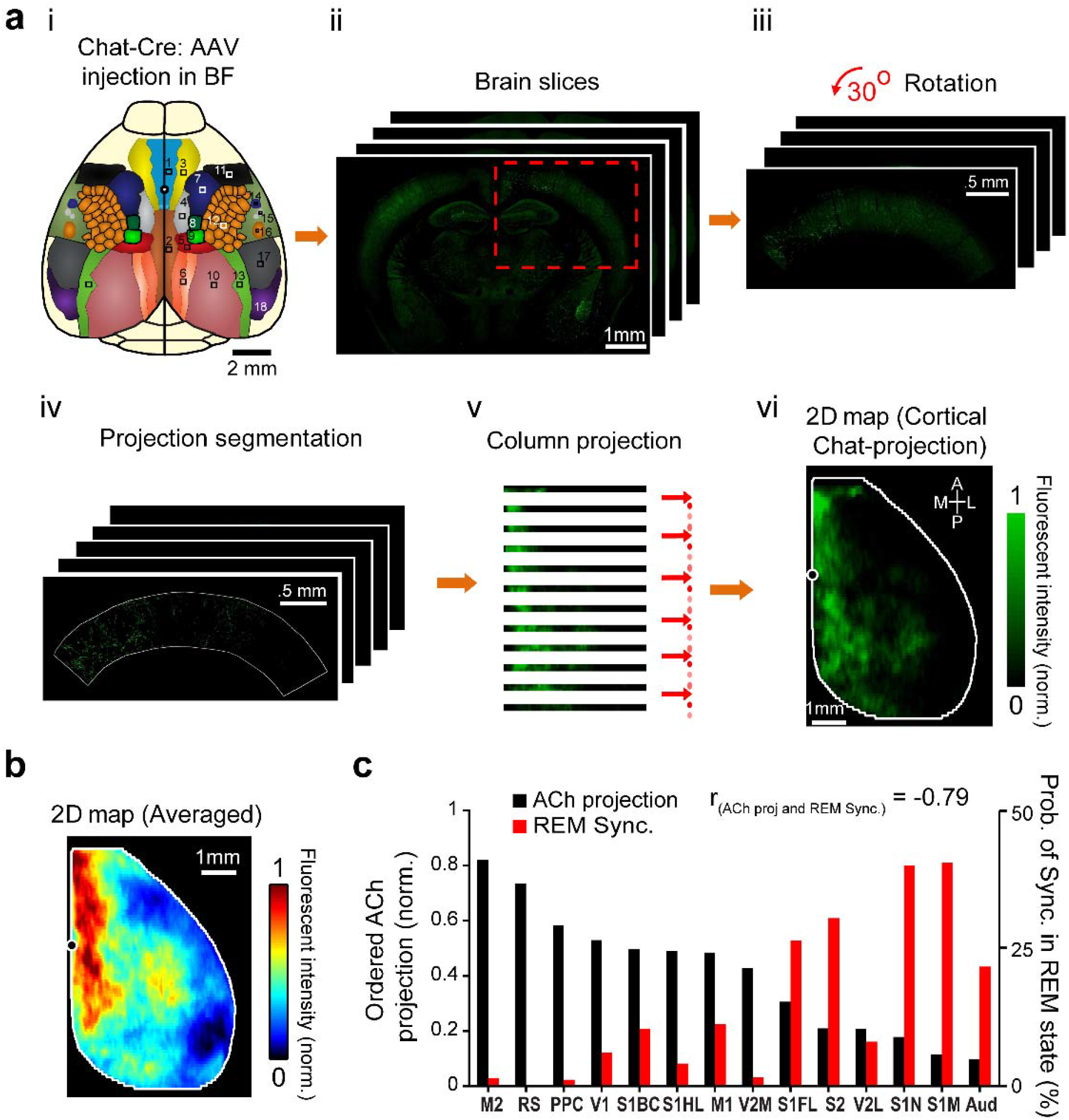
The strength of cortical cholinergic axonal projections from the basal forebrain (BF) is negatively correlated with the probability of SWA during REM sleep. (a) (i) A Cre-dependent AAV expressing anterograde GFP tracer was injected into the distinct non-overlapping BF nuclei of Chat-IRES-Cre mice to map BF cholinergic projections to different cortical regions. To quantitatively assess the cholinergic innervation of the cortex, 2D representations of Allen Mouse Brain Connectivity (AMBC) data sets were created for the dorsal cortex (ii–vi, see methods). vi, Note higher distribution of cholinergic innervation within anterior-posterior axis of midline cortical regions. The map is obtained from a single BF injection. (b) Averaged cholinergic innervation map from 10 injection BF sites. (c) Comparison of the strength of BF cholinergic projections to different ROIs (black, n=10) with the probability of synchronized activity during REM state (red, combined glutamate and VSD data, n=13). ROIs are sorted based on the strength of the cholinergic innervation they receive. The strength of ACh projections was negatively correlated with the synchronization probability (r=−0.79, p<0.001).

It is known that VSD and glutamate signals mainly measure activity from superficial cortical layers; however, the BF cholinergic map, discussed above, was calculated based on projections to all cortical layers. To address this caveat, we compared the BF cholinergic projections to the supragranular layers I-III with those to all cortical layers and found that the cholinergic projection maps were similar (supplementary Fig. 8a, r=0.78, p<0.001).

To determine if increasing the ACh level could affect the local SWA during REM sleep, six unrestrained mice were intraperitoneally (IP) injected with donepezil (1 mg/kg), a cholinesterase inhibitor that increases the acetylcholine level globally in the brain. We found that donepezil significantly reduced the probability of synchronized activity in REM sleep (supplementary Fig. 8b). Amphetamine, a brain stimulant drug, is known to desynchronize cortical activity by affecting multiple neuromodulatory systems. It increases acetylcholine, noradrenaline, dopamine, and glutamate levels, and reduces extracellular GABA concentrations^39–41^. In three animals, after performing VSD imaging under urethane, we systemically induced a REM-like state by IP injection of amphetamine (0.1 mg/kg). This injection induced slow theta (3.5±0.25 Hz; n=3 mice) activity in the hippocampus, and globally desynchronized activity in the cortex for at least 30 min (with an onset latency of 15-30 min)^42^ (supplementary Fig. 9a). In other words, amphetamine abolished synchronized activity in all cortical regions during the REM-like state (supplementary Fig. 9b).

The variations in cholinergic axonal projections would not necessarily correlate with those in terminal activity. To shed light on the functional role of acetylcholine (ACh) in local desynchronization, we recorded the activity of cholinergic terminals in the cortex during REM sleep. Ai162^43^ and Chat-cre mice were crossed in order to express the calcium sensor GCaMP6s in cholinergic neurons and their axonal projections (Fig. 4a and supplementary Fig. 10a,b). These mice were then used to infer fluctuations in cholinergic tone across NREM and REM sleep. ACh activity during sleep was assessed by combining widefield calcium imaging of cortical cholinergic activity with hippocampal LFP recording. The data was recorded in one ~1.5 hour session, during which several NREM to REM sleep transitions occurred.

Spontaneous transition of the forebrain from NREM to REM sleep was associated with a general increase in the calcium signal (after detrending), consistent with an increase of the cholinergic axonal and neuronal activity in the cortex (Fig. 5b,c). The increase in cholinergic activity during REM sleep was not homogeneous across all cortical regions (Fig. 5e,f).

**Figure 5.**
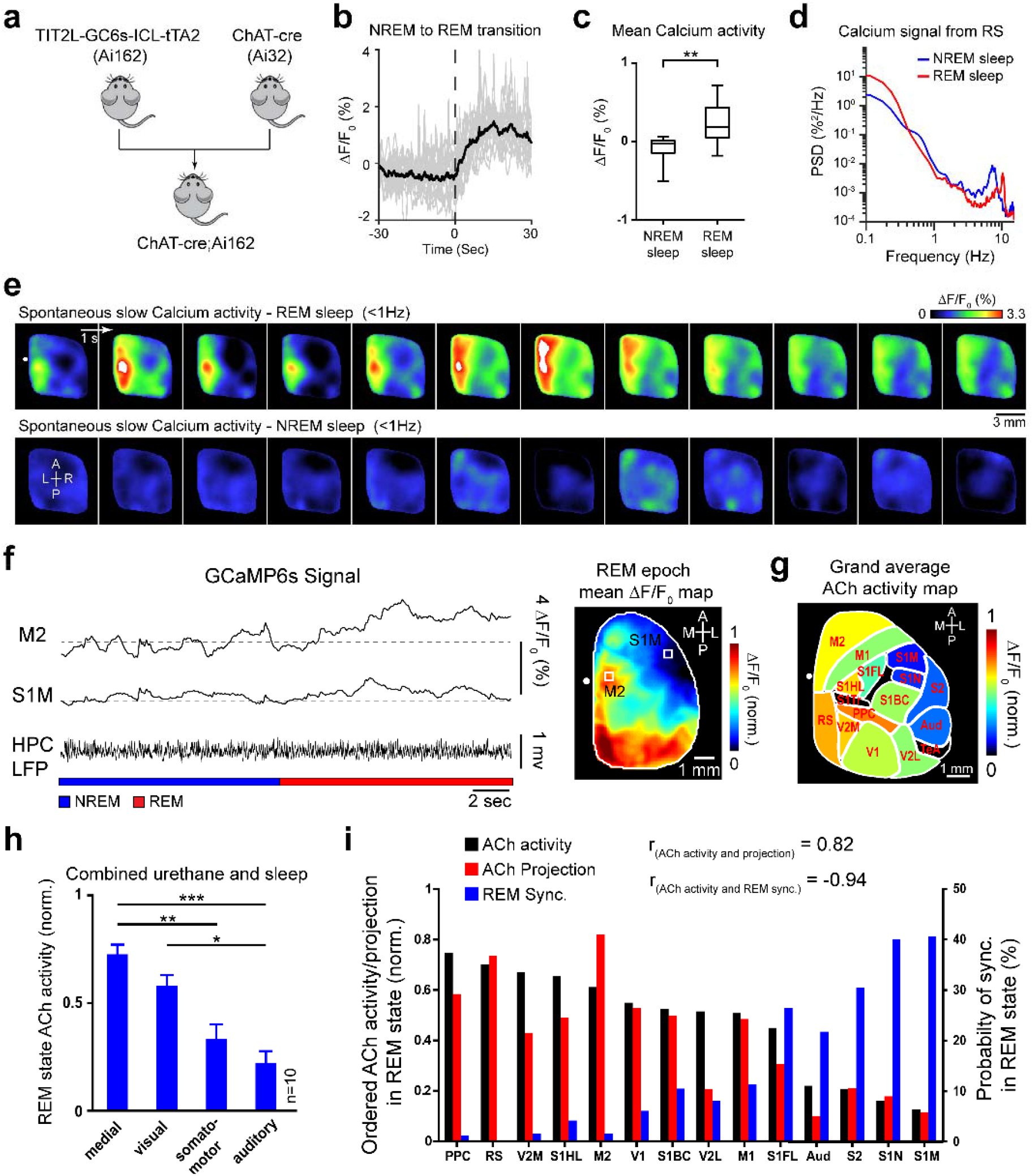
The level of cortical cholinergic activity captured by calcium imaging is negatively correlated with the probability of SWA during REM sleep. (a) Ai162 and ChAT-cre transgenic lines (Jackson Lab) were crossed to produce mice in which GCaMP6s sensor is specifically expressed in cholinergic neurons. (b) Change of calcium fluorescence (∆F/F_0_ averaged over entire cortex) during NREM to REM sleep transition. Gray and black lines indicate individual and average transitions respectively. Dashed line indicates the transition onset from NREM to REM sleep. (c) Boxplot of episode-wise mean calcium fluorescence during NREM and REM sleep (n=16 REM episodes in four mice, paired t-test, ** p<0.01). (d) Power spectral density of calcium signal from retrosplenial cortex for two representative episodes of NREM and REM sleep. (e) Montages show spatiotemporal dynamics of calcium activity measured from cholinergic fibers and neurons during REM-like and NREM-like states. (f) Left, changes in the Ca^2+^ signal measured from two cortical regions (S1M and M2) during the transition from a NREM sleep to a REM sleep episode. Dashed lines show baseline fluorescence (F_0_). Right, map of mean calcium activity calculated for the representative REM sleep episode in left. (g) Similar to f-right, but animal-wise average map (n=10 animals) registered onto the Allen Institute Mouse Brain Coordinate Atlas. Note that lateral regions show lower level of ACh release, indirectly measured by Ca^2+^ imaging. (h) Animal-wise average (n=10) of ACh activity across cortical subnetworks, sorted in descending order. Error bars, s.e.m. (repeated measure ANOVA with Greenhouse-Geisser correction for sphericity: F3,27 = 17.481, p = 1.658×10^−6^; post-hoc multiple comparison with Tuckey’s correction: medial vs visual p = 0.123, medial vs auditory p = 8.04×10^−4^, medial vs somatomotor p =1.66×10^−3^, visual vs auditory p = 0.0153, visual vs somatomotor p = 0.094, auditory vs somatomotor p = 0.36). (i) Ordered level of ACh activity measured in 14 ROIs (black, n=10) compared to the strength of BF cholinergic projections to the same ROIs (red, n=10) and the probability of synchronized activity during REM state (blue, n=13). ROIs are ordered based on their ACh activity during REM state. (p<0.001 for both Spearman’s rank correlations).

Furthermore, the spectral analysis of the calcium signal revealed that most of the power of cholinergic terminal activity was concentrated in the slow (<1 Hz) frequency band (Fig. 5d).

We also managed to record brain activity in another group (n=6) of these ChAT□JCre;Ai162 mice under urethane anesthesia. We found that ACh terminal activity in the REM-like state of urethane anesthesia is similar to REM sleep in the medial, visual, somatomotor, and auditory subnetworks (supplementary Fig. 10c). Based on this, we combined the natural sleep and urethane data. For 10 mice (4 sleep + 6 urethane), we registered 14 cortical regions on the map of average cholinergic activity based on anatomical landmarks and we grouped these regions into 4 subnetworks (Fig. 5g). Medial and visual subnetworks had the highest level of ACh activity, followed by somatomotor and auditory networks (Fig. 5h; n = 10 mice per subnetwork; repeated measure ANOVA with Greenhouse-Geisser correction for sphericity: F3,27 = 17.481, p = 1.658×10^−6^; post-hoc multiple comparison with Tuckey’s correction: medial vs visual p = 0.123, medial vs auditory p = 8.04×10^−4^, medial vs somatomotor p =1.66×10^−3^, visual vs auditory p = 0.0153, visual vs somatomotor p = 0.094, auditory vs somatomotor p = 0.36). Moreover, the level of cholinergic activity across cortical regions showed a significant positive correlation with the strength of BF cholinergic projections to these regions, and also a significant negative correlation with the probability of synchronized activity during the REM state (Fig. 5i; r_ACh activity and projection_=0.82, r_ACh activity and REM sync._ =−0.94, p < 0.001 for both correlations).

Moreover, we mapped the patterns of cholinergic sensory-evoked activity using calcium imaging in urethane anesthetized mice. All modes of sensory stimulation (auditory, visual, whisker, and hindlimb) resulted in activation of cholinergic terminals in the corresponding primary sensory cortical area, followed by a spread of activity toward secondary and functionally related cortical regions (supplementary Fig. 10d). This was in line with modality-specific patterns of activation recorded using voltage imaging^25^ and points to the recruitment of acetylcholine in the processing of sensory inputs.

We also performed control experiments to rule out the possibility of non-neuronal processes, including hemodynamic response, contaminating our calcium imaging data. For that, we incubated the exposed portion of the brain with lidocaine, and it completely abolished the response to hindlimb stimulation in all cortical regions (supplementary Fig. 10e, f). We next examined the effect of lidocaine administration on the spontaneous ACh fluctuations recorded with calcium imaging. To do so, we calculated the root mean square (RMS) of calcium activity at each pixel over a 20 min window before and after lidocaine application. Based on the resulting maps, RMS values reduced throughout the cortex, and the pixel-wise average of spontaneous activity RMS decreased following lidocaine administration (supplementary Fig. 10g, h).

## Discussion

The regional variations in cortical desynchronization during REM sleep and the REM-like state were assessed using electrophysiology and various wide-field optical imaging techniques in sleeping and urethane anesthetized mice. In head-fixed sleeping mice, glutamate imaging revealed that, even if the hippocampal LFP signal clearly indicated that the brain was in REM sleep, there were often certain cortical regions that continued to exhibit different gradient of slow-wave activity, which is generally characteristic of NREM sleep. We identified spatial gradient of slow-wave activity during REM centered around the sensorimotor and auditory cortical subnetworks, including the forelimb, mouth and nose primary somatosensory, secondary somatosensory, and auditory cortices. This network, both in our data and in the anatomical characterization is orthogonal with the Default Mode Network in the mouse brain^13,22^.We further investigated whether local synchronization patterns could be detected during REM sleep using two independent approaches: electrophysiological recordings from freely behaving mice, and voltage imaging of the cortex in urethane anesthetized animals. These experiments confirmed that synchronized patterns did occur in localized regions of the cortex during both REM sleep and the REM-like state. Furthermore, the spatial topography of local activity patterns was very similar to that observed with glutamate imaging, with the somatomotor and auditory cortical regions being more likely to show synchronized activity during the REM state.

The results are consistent with a previous electrophysiological study in mice^9^ reporting that the primary somatosensory, motor, and visual areas show low-frequency oscillations in REM sleep, while retrosplenial and secondary motor/visual cortices demonstrate desynchronized patterns. Our results are also in accordance with a human EEG study which reported increased low-frequency oscillations in primary somatosensory, motor, and visual cortices during REM sleep compared to wakefulness^12^. Similar to our data, in these two studies, the primary visual cortex exhibited weak, low-frequency activity during REM sleep.

In the present study, we also investigated the mechanisms underlying local desynchronization in REM sleep. Cholinergic tone is well-known to increase during REM sleep; however, it is also known that cholinergic innervation varies over the cortex^36,37^, as do many indicators of cholinergic neurotransmission such as AChE, ChAT, muscarinic, and nicotinic receptors^44–48^. These studies demonstrated that the entorhinal and olfactory cortices tend to be more densely innervated by cholinergic axons, yet they did not conduct systematic measurement of cholinergic axons in all the cortical regions that were imaged in the present study. We hypothesized that regions of the cortex that receive less cholinergic innervation from the basal forebrain would be more likely to show slow-wave activity during REM sleep, relative to cortical regions with more abundant cholinergic innervation (and, presumably, greater levels of acetylcholine release). To test this, we used Allen Mouse Brain Connectivity (AMBC) data to measure the density of cholinergic projections from the basal forebrain to different cortical regions. In support of our hypothesis, we found that the amount of cholinergic innervation was negatively correlated with the likelihood of a cortical region exhibiting synchronized activity during REM sleep. The midline cortical areas, which were most likely to transition to REM sleep, receive the strongest cholinergic innervation from the basal forebrain.

To explore whether cholinergic neural activity per se underlies the variations in cortical desynchronization during REM sleep, we imaged the activity of cholinergic fibers throughout the cortex using calcium imaging. We confirmed that the somatomotor and auditory cortical regions, the epicenters of synchronized activity in REM sleep in our findings, have lower levels of cholinergic activity during REM sleep. Furthermore, by injecting donepezil, which blocks the breakdown of acetylcholine, we demonstrated that increasing the acetylcholine level significantly reduces the probability of synchronization in REM sleep. These results suggest that acetylcholine is one of the key players for the local cortical state alternation in REM sleep. Our results are consistent with another study in mice, which showed that ACh release during whisking suppresses slow-wave activity in the sensory cortex^49^. To the best of our knowledge, this is the first study to image cholinergic activity across a wide area of the mouse cortex during sleep. Here, the calcium signal reflects the activity of cholinergic axons stemming from the basal forebrain as well as the activity of cortical cholinergic interneurons. The interneurons account for between 12% and 30% of cortical cholinergic signaling, and indirectly increase excitably in neighboring neurons^50,51^. Dissecting the role of these local cholinergic interneurons from basal forebrain cholinergic projections is one possible direction for future studies. Another caveat of our approach in investigating the cholinergic system is that it overlooks other possible factors, including differences in the patterns of cholinergic receptor expression and possible differences in the efficacy of acetylcholine degradation and reuptake throughout the cortex.

Besides acetylcholine, other neuromodulatory systems can influence the pattern of cortical synchronization and desynchronization during REM sleep. It has previously been shown that noradrenergic projections, originating mainly from the locus coeruleus, can desynchronize multiple sensory cortices^34^. Consistent with this, we provide evidence that increasing noradrenaline levels by systemic administration of amphetamine globally desynchronizes the cortex and abolishes slow-wave activity during the REM state. Therefore, it is likely that low levels of noradrenaline during REM sleep are required for the occurrence of local cortical synchronized patterns.

Although it is still not clear how NREM and REM sleep contribute to brain plasticity, it is tempting to interpret our findings by the sequential hypothesis^52,53^, which holds that NREM and REM sleep manipulate brain circuits differently and sequentially. During NREM sleep, plasticity is reduced, and the brain often reactivates cell assemblies that correspond to recent experiences (i.e., memory reactivation). During the subsequent REM sleep, plasticity is increased, and modification of synapses that were potentiated within the reactivated cell assemblies takes place^53^. Therefore, according to the hypothesis, it is important to have episodes of both NREM and REM sleep sequentially. Our results show that sequential alternation between NREM and REM sleep is less common in the anterior and lateral cortical areas relative to the midline and occipital cortical areas. This may indicate that synaptic modification is slower or suppressed in the anterior and lateral compared with the midline and occipital cortical areas. Many of the anterior and lateral areas in our experiment consisted of primary and secondary sensory areas. These areas may have more connections that were genetically prewired than the association areas along the midline. The disrupted NREM-REM sleep sequences in the anterior and lateral areas may help preserve the prewired connections.

Additionally, our lab and others have shown that hippocampal sharp-wave ripples, the main candidate events for memory reactivation during NREM sleep, are well correlated with activation of default mode network which includes midline and occipital cortical areas^14,15^. This coordination is followed by a strong cholinergic activity and desynchronized cortical patterns, which enhance synaptic plasticity^53–55^ in these regions during subsequent REM sleep. This can ultimately serve the synaptic consolidation and fulfill the sequential sleep hypothesis.

## Methods

### Animals

A total of 23 adult C57BL/6J mice, male and female, aged 2-4 months and weighing ~30 g, were used for imaging experiments. For head-fixed glutamate imaging, 6 adult EMX-CaMKII-iGluSnFR transgenic mice expressing iGluSnFR in glutamatergic cortical neurons were used. Seven C57BL/6J mice were used for acute voltage-sensitive dye imaging under urethane anesthesia. For ACh imaging study, Ai162 and Chat-cre mice were crossed in order to express the calcium sensor GCaMP6s in cholinergic neurons (6 and 4 mice for urethane and sleep experiments respectively). An additional set of 8 C57BL/6J mice were used for electrophysiological study. All animals were housed under 12:12 h dark/light cycle and had ad libitum access to water and food at all time. Animal protocols were approved by the University of Lethbridge Animal Care Committee and followed the guidelines issued by the Canadian Council on Animal Care.

### Acute imaging preparation

#### Surgery

Mice were anesthetized with urethane (0.12% wt/wt), and a large unilateral craniotomy (6.5 × 6 mm; bregma 2.8 to −3.7 mm and lateral 0 to 6 mm) was made. Underlying dura matter was removed. Body temperature was kept at 37 ± 0.5 °C using a feedback loop heating pad. To assist with breathing, a tracheotomy was performed. Additional doses of urethane (10% of initial dose) were administered when necessary to keep the surgical plane of anesthesia.

#### Voltage-sensitive dye imaging

RH1961 (Optical Imaging) was dissolved in HEPES-buffered saline solution (0.5 mg/ml) and applied for 40-60 min to the cortex exposed by the craniotomy. Unbounded dye molecules were washed away afterward. For minimizing respiration artifacts, 1.5% agarose made with HEPES-buffered saline was spread over the cortex and sealed by a coverslip. VSD image stacks were collected in 12-bit format at 100 Hz frame rate using a CCD camera (1M60 Pantera, Dalsa, Waterloo, ON) and an EPIX E8 frame grabber controlled with XCAP 3.7 imaging software (EPIX, Inc, Buffalo Grove, IL) (1M60 Pantera, Dalsa) and XCAP 3.8 imaging software (EPIX, Inc.). Images were taken through a microscope composed of front-to-front video lenses (8.6 × 8.6 mm field of view, 67 μm per pixel). VSD was excited by two red LEDs (627-nm center, Luxeon K2) and emitted fluorescence passed through a 673-nm to 703-nm bandpass emission filter. Each VSD imaging stack of spontaneous activity included 90,000 frames.

#### Electrophysiological recording

The hippocampal electrode was implanted outside the planed cranial window prior to performing the craniotomy. For that, a Teflon-coated stainless steel wire (bare diameter 50.8 μm) was placed in the pyramidal layer of the right dorsal hippocampus. It was inserted posterior to the occipital suture with 33-degree angle (with respect to the vertical axis) according to the following coordinates relative to bregma: mediolateral (ML): 2.3 mm; dorsoventral (DV): 1.6 to 1.9 mm. The position of the electrode tip was confirmed using an audio monitor (Grass Instrument Co.). For cortical recording, a bipolar electrode made with Formvar-coated nichrome wire (coated diameter 38 μm) was inserted into either the primary motor or auditory sensory cortex; the tip separation was 0.5 mm and the upper tip was located in layer 2/3. The reference and ground electrodes were placed on the cerebellum. The LFP and EMG signals were amplified (×1000) and filtered (0.1-10,000 Hz) using a Grass P5 Series AC amplifier (Grass Instrument Co.) and were sampled at 20 kHz using a data acquisition system (Axon Instruments). After data collection, 100 μA current was injected into the hippocampal electrode for 10 sec. Animals were sacrificed and brains were extracted, sectioned and mounted. Location of the hippocampal electrode was further confirmed using cresyl violet staining.

#### Amphetamine administration

After recording 4-5 imaging stacks of spontaneous activity, animals were injected with amphetamine (IP, 0.1 mg/kg). After 20-30 min, the effect of the drug on brain activity could be detected based on the ongoing electrophysiological recording. From here, electrophysiological recording and VSD imaging were resumed.

### Chronic imaging preparation

#### Surgery

Six mice were anesthetized with isoflurane (2.5% induction, 1-1.5% maintenance) and, using antiseptic techniques, were implanted with hippocampal and EMG electrodes. For the hippocampal electrode, a bipolar electrode made from a Teflon-coated stainless-steel wire (bare diameter 50.8 μm, tip separation 0.5 mm) was placed in the pyramidal layer of the right dorsal hippocampus. It was inserted posterior to the occipital suture with 33-degree angle (with respect to the vertical axis) according to the following coordinates relative to bregma: ML: 2.3 mm; DV (upper tip): 1.6 to 1.9 mm. For EMG recording, a bipolar multi-stranded stainless-steel wire (bare diameter 127 μm) was inserted into the neck musculature using a 22-gauge needle. The ground screws were placed on the skull over the cerebellum. Then the skull was covered with a thin layer of transparent metabond (Parkell, Inc.), and a headplate was secured to it using metobond. The other ends of the hippocampal and EMG electrodes were soldered to a connector (Mill-Max Mfg. Corp.), and the connector was fixed on the edge of the headplate.

#### Habituation and sleep restriction

Animals were allowed to recover for 7 days after surgery and were then habituated to the recording setup for at least 2 weeks. For habituation, animals were moved to the recording platform in the morning and were head-restrained using two clamps. Their body motion was limited by putting them in a plastic tube. Sounds and vibrations in the recording setup and the room were minimized, and conditions of the recording day were simulated as much as possible. In the first session, mice were allowed to explore the platform without restraint. In the next session, mice were kept head-fixed for 5 min, and this time was increased by 5-10 min every day over the next 2 weeks. After each session, the animals were rewarded with two Cheerios.

Sleep restriction was performed the day before the recording session. To do so, animals’ home cages were transferred from the housing room to a procedure room around 2 pm. There, the animal’s sleep was restricted by mild external stimulation using a cotton applicator for 1-2 seconds whenever drowsiness (immobility with eyelid closure) was observed. We continued the sleep restriction for 6-8 hours and then transferred the animals to a cage with a few objects and a running wheel. The animals spent the night in this cage and had ad libitum access to food and water. In the morning around 8 AM, animals were transferred to the recording setup for combined glutamate and electrophysiological recording. After the recording session, animals were returned to their home cage and to the housing room where they could sleep undisturbed for the rest of the day. They were allowed to recover for a week before the next recording session.

#### Glutamate imaging

Animals were gently moved to the recording setup and head-fixed using two clamps. Room temperature was maintained at 25 °C, and the recording platform was kept warm using a microwavable heating pad. Part of the animal’s nesting materials were put close to them to reduce stress. For exciting the iGluSnFR, two blue LEDs (470-nm center, Luxeon K2) were used. They were turned on at the beginning of the recording and kept on throughout the imaging session. Image stacks were collected at 100 Hz frame rate. Emitted fluorescence passed through a 510-nm to 550-nm bandpass emission filter. Three stacks of spontaneous activity (170,000 frames each) were recorded during each session. Recordings were repeated once a week.

#### Electrophysiological and video recording

The hippocampal LFP and EMG signals were amplified (×1000) and filtered (0.1-10,000 Hz) using a Grass P5 Series AC amplifier (Grass Instrument Co.) and were sampled at 20 kHz using a data acquisition system (Axon Instruments). The animal’s behavior and pupil movements were filmed at 30 Hz using a V2 infrared Pi camera. The animal’s head, shoulders, and forelimbs were illuminated using a 940 nm infrared LED.

#### Calcium Imaging of cholinergic terminal activity

Procedures for hippocampal electrode implantation and chronic window preparation were similar to glutamate imaging, except the mice had GCaMP6s indicators expressed in their cholinergic cells. Animals were habituated for two weeks, and sleep restricted the day before recording, as explained before. Image stacks were collected for calcium at 30 Hz frame rate. GCaMP6s indicators were excited by two blue LEDs (470-nm center, Luxeon K2), and emitted fluorescence passed through a 510-nm to 550-nm bandpass emission filter. Each imaging stack of spontaneous activity included 170,000 frames.

### Chronic EEG recording

#### Surgery

Seven C57BL/6J mice were anesthetized with isoflurane (2.5% induction, 1-1.5% maintenance) and implanted with cortical, hippocampal, and muscular electrodes, using aseptic techniques. For cortical and hippocampal electrodes, bipolar (tip separation =.6 mm) and monopolar electrodes made from Teflon-coated stainless steel wire (bare diameter 50.8 μm) were implanted in cortical areas, and in the pyramidal layer of the CA1 hippocampal region according to the following coordinates (in mm): primary motor cortex (M1): AP: 1.5, ML: −1.7, DV: 1.5, secondary motor cortex (M2): AP: 1.7, ML: 0.6, DV:1.1 mm, mouth primary somatosensory area (S1M): AP: 0.85, ML: 2.8, DV: 1.4, barrel primary somatosensory area (S1BC): AP: −0.1, ML: −3.0, DV: 1.4, retrosplenial cortex (RS): AP: −2.5, ML: 0.6, DV: 1.1, and hippocampus (HPC): AP: −2.5, ML: 2.0, DV:1.1 mm. For EMG, a multistranded Teflon-coated stainless-steel wire (bare diameter 127 μm) was implanted into the neck musculature using a 22 gauge needle. The reference and ground screws were placed on the skull over the cerebellum. The other ends of the electrode wires were clamped between two receptacle connectors (Mill-Max Mfg. Corp.), and the headpiece was secured to the skull using metabond and dental cement.

#### Electrophysiology

Animals were allowed to recover for 7 days after surgery, and then habituated for 5-7 days in the recording setup. On baseline days, animals were injected with saline at 8:25 AM and moved to the recording setup, where their sleep activity was recorded from 8:30 AM for 4 hours using a motorized commutator (NeuroTek Inc.). Baseline recording was repeated for 3 days. On the fourth day, animals were injected with donepezil (IP, dosage 1 mg/kg) 5 minutes before the recording. LFP and EMG recordings were amplified, filtered (0.1-4000 Hz), and digitized at 16 kHz using a Digital Lynx SX Electrophysiology System (Neuralynx, Inc.), and the data was recorded and stored on a local PC using Cheetah software (Neuralynx, Inc.). A Pi camera was used to record the animal’s behaviour during the recording.

### Data analysis

#### State scoring in urethane anesthesia

LFP and EMG signals were downsampled to 2 kHz offline, and analyzed using custom-written codes in MATLAB (MathWorks). REM-like, NREM-like, and transition states were scored based on the hippocampal LFP. The ratios of theta (3-5 Hz) power and slow wave (0.2-1.2 Hz) power to the total power were calculated in 4-sec windows, and a threshold of mean plus one standard deviation was used to differentiate the states. The hippocampal LFP was scored as a REM-like state when the theta ratio was above the threshold for 10 consecutive seconds while the slow wave ratio was below the threshold simultaneously. A NREM-like state was defined as an opposite case of the REM-like state; that is, when the slow wave ratio was above the threshold for 10 consecutive seconds while the theta ratio was below the threshold simultaneously. A period that did not meet these conditions was classified as a transition state.

#### Quantification of pupil diameter

Pupil diameter during different states of sleep and wakefulness was quantified offline using a custom algorithm implemented in Bonsai (https://bonsai-rx.org//). Briefly, the algorithm detects the pupil by segmenting it from the iris and sclera and models it as an ellipse. The major axis length of this ellipse was quantified as the pupil diameter.

#### Sleep scoring

LFP and EMG signals were downsampled to 2 kHz offline. Scoring was done in 4-sec epochs. The ratios of theta (5-10 Hz) power and delta (0.5-4 Hz) power to the total power were calculated for the hippocampal LFP signal in 4-sec windows. Raw EMG activity was filtered between 70 Hz and 1000 Hz and then rectified and integrated using 4-sec moving windows. A state was scored as NREM sleep when the EMG signal was smaller than half of its mean and the delta power ratio was above 0.4-0.5. REM sleep was detected when the EMG signal was smaller than half of its mean and the theta power ratio was above 0.4-0.5. To avoid the transitory periods, the first 10 sec and last 5 sec of the REM episodes were excluded. Everything else was scored as wakefulness. This scoring was confirmed by another observer separately. In the head-fixed recording preparation, the pupil diameter of the animals was used to assist the observer in correcting the scoring; However, when there was contradiction between scoring based on pupil diameter and electrophysiological recording, the latter was used as criteria.

For scoring phasic versus tonic REM sleep in glutamate mice, face movement was detected using a camera that recorded animal and pupil movement. To define animal movement, a 25×25-pixel square was selected on the animal’s chin in the behavioral video, and the average intensity of this ROI was calculated over time. This signal was filtered above 1 Hz and thresholded using the mean + 1 SD to detect chin movement. Each 3-sec epoch which had movement in more than 20% of it was considered to be a phasic REM event.

#### Image preprocessing

Raw data for VSD and glutamate imaging of spontaneous activity were preprocessed based on the following steps: First, the time series of each pixel was filtered using a zero-phase highpass Chebyshev filter above 0.2 Hz. Then, a baseline signal (F_0_) was calculated by averaging all the frames, and the fluorescence changes were quantified as (F − F_0_)/ F_0_ × 100, where F − F_0_ is the filtered signal. PCA was subsequently performed on the image stacks, and the principal components with the greatest singular values were kept. To further reduce spatial noise, images were filtered by a Gaussian kernel (5 × 5 pixels, sigma = 1). To eliminate the strong heartbeat artifact, imaging data was also filtered using a zero-phase lowpass Chebyshev filter below 6 Hz. In calcium imaging data, ∆F was calculated using the locdetrend function in the Choronux toolbox, in which a piecewise linear curve was fitted to the pixel time series using the local regression method (150-sec moving window, 100-sec step size). The rest of the steps were similar to glutamate imaging.

#### Calculation of p-value maps of synchronization

P-value maps were calculated by comparing REM and NREM sleep SW/delta powers. For that, imaging stacks of each NREM episode (defined based on hippocampal LFP and EMG) from one continuous recording session were first concatenated in time. Then the 0.5 to 5-Hz power (multitaper method) of the optical signal for each pixel was calculated in 3-sec long windows (2-sec overlap). Same was repeated for each 8-sec epoch of REM sleep. This generates two power distributions for the concatenated NREM data and the individual REM epochs (for each pixel). To determine if a specific pixel during a given REM epoch undergoes state transition, a test of significance (Wilcoxon rank-sum test, α = 0.05) was performed against the null hypothesis that the median of the REM distribution is equal to that of the NREM distribution. The pixel activity was scored as desynchronized if the calculated p-value was significant, otherwise it was scored as synchronized.

#### Calculation of p-value map overlap

P-value maps calculated for each REM sleep epoch were first binarized based on 0.05 significance level (1 was assigned to pixels that had p-value greater than and equal to 0.05, and 0 was assigned to the rest). Only maps that contained at least 100 nonzero pixels were kept. Overlap between each of the two maps was calculated based on this formula:

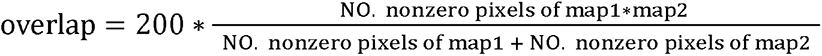

#### Calculation of two-dimensional map of cortical cholinergic projection

Images of coronal sections of mice brain injected with GFP-expressing AAV in the basal forebrain nuclei were downloaded from Allen Institute for Brain Science (http://connectivity.brain-map.org/). Each section was 100 μm apart and its resolution was 2.8 μm per pixel. For each injection experiment, a series of coronal section images spanning the entire cortex was assembled into a 3D stack. This stack was rotated laterally 30° to match the angle of the mouse head in the glutamate and VSD imaging experiments. Projection data outside of the cortex was masked to remove viral expression in subcortical areas. The 3D data matrix was converted to a 2D matrix by averaging the entries across the dimension associated with cortical layers. 2D maps obtained from different injection experiments were registered using midline and position of bregma and then averaged together.

#### Statistical tests

All statistical test for linear data were performed using MATLAB built-in functions. Paired t-test and Wilcoxon signed-rank tests were used for linear data. Error bars and ± ranges represent the s.e.m. *P < 0.05, **P < 0.01, ***P < 0.001. All the statistical tests were two sided.

## Author contributions

Conceptualization, M.N., M.H.M.; Methodology, M.N., J.K.A., M.Nagh., E.J.B.C.; Investigation, M.N., M.H.M., M.T., B.L.M.; Formal Analysis, M.N.; Writing – Original Draft, M.N., M.H.M.; Writing – Review & Editing, M.N., M.H.M., M.T., B.L.M., J.K.A., M.Nagh., E.J.B.C.; Funding Acquisition, M.H.M.; Resources, M.H.M.; Supervision, M.H.M.

## Acknowledgements

This work was supported by Natural Sciences and Engineering Research Council of Canada (grant no 40352 & 1631465 to MHM and BLM respectively), Alberta Innovates (MHM and BLM), Alberta Prion Research Institute (grant no. 43568 to MHM), and Canadian Institute for Health Research (grant no 390930 & 156040 to MHM and BLM MHM respectively), National Science Foundation (MHM, BLM), and USA Defense Advanced Research Projects Agency (grant no HR0011-18-2-0021 to BLM). The authors thank Di Shao and Behroo Mirza Agha for animal breeding, Jianjun Sun for surgical assistance, Brendan McAllister and Emily Hagens for revision of the manuscript. The authors thank Hongkoi Zeng from the Allen Institute for Brain Science for providing the Emx-cre, Camk2a-tTa, and Ai85 mice as a gift.

## Declaration of Interests

The authors declare no competing interests.

---

**Supplementary Figure 1.**
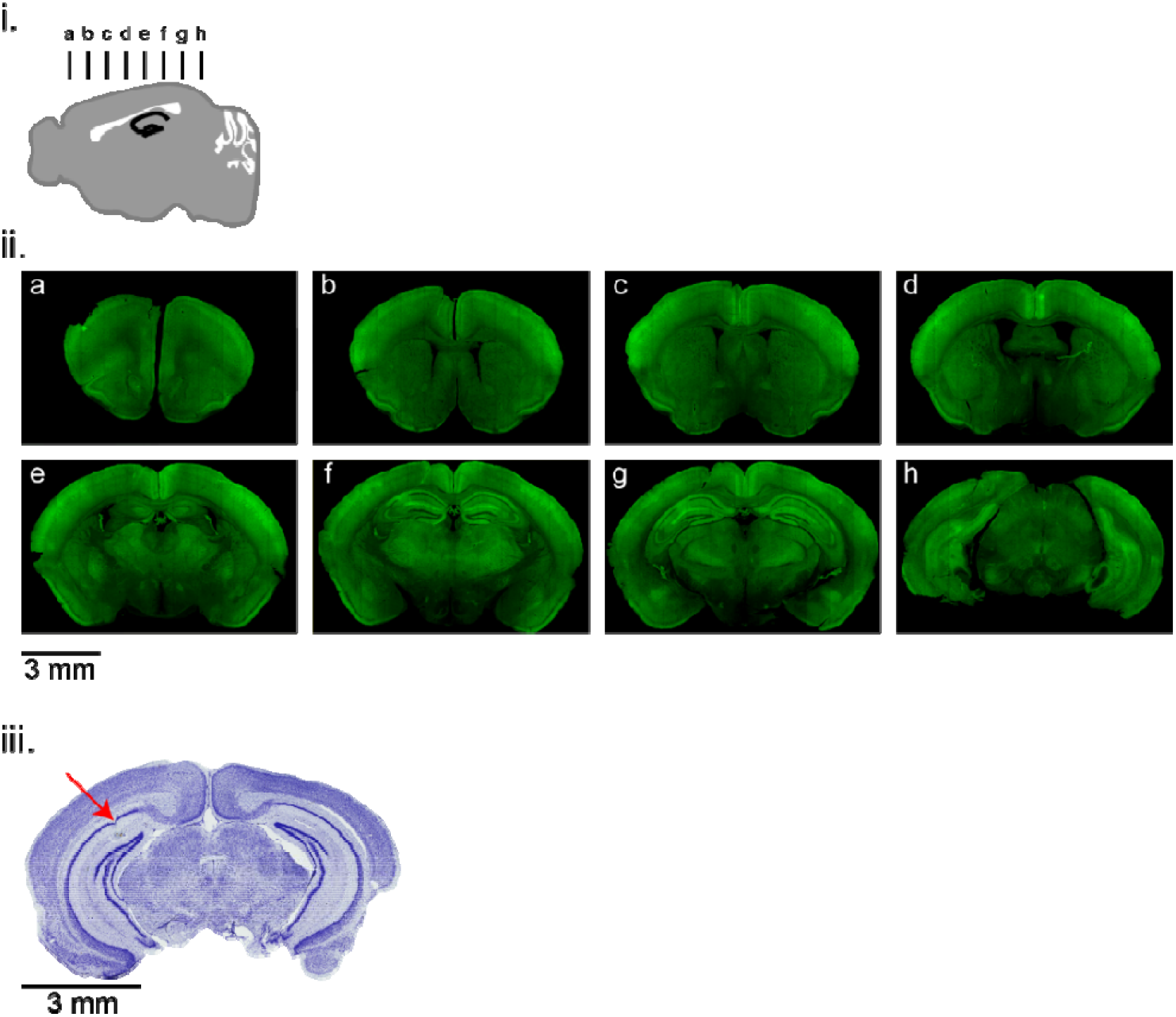
Expression of iGluSnFR and electrode localization. (i) Schematic diagram showing the location of brain sections in panel ii. (ii) Coronal brain sections show the expression of iGluSnFR in neocortex and hippocampus in Ai85-CamKII-Emx mouse. (iii) Representative brain section stained with nissle, showing the tip location of the hippocampal electrode.

**Supplementary Figure 2.**
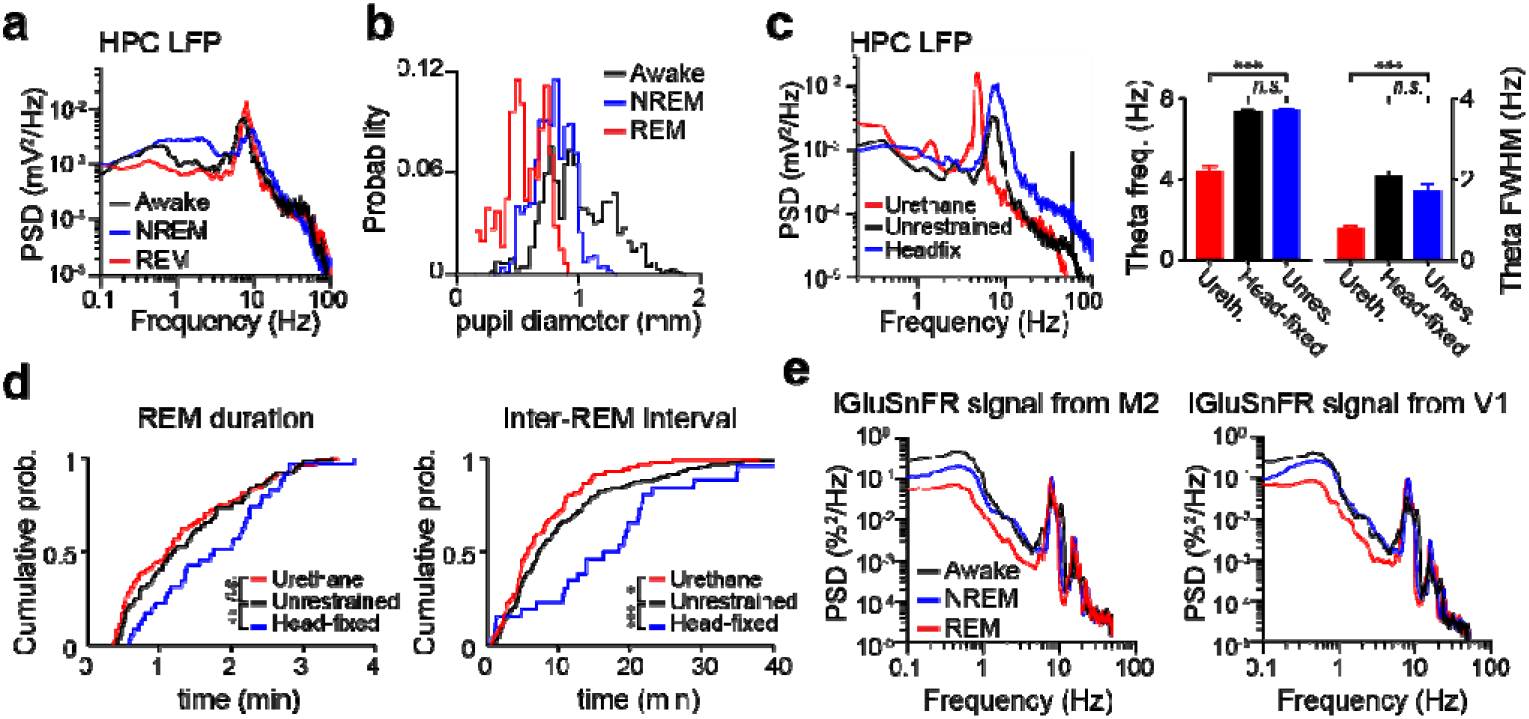
(a) Power spectral density (PSD) of hippocampal LFP during waking and NREM/REM sleep. (b) Distribution of pupil diameter across behavioral states in a representative recording session (c) Comparison of hippocampal theta activity (left: PSD, right: theta peak frequency and full width at half maximum (FWHM) frequency between head-fixed REM sleep (n=6), unrestrained REM sleep (n=8), and urethane REM-like state (n=7). Error bars, s.e.m. (***P < 0.001, Student’s t-test). (d) Cumulative distribution of REM and inter-REM bouts duration in head-fixed, unrestrained and anesthetized mice (**P < 0.01, ***P < 0.001, Kolmogorov-Smirnov test). (e) Power spectral density of iGluSnFR signal from M2 (left) and V1 (right) during waking and NREM/REM sleep.

**Supplementary Figure 3.**
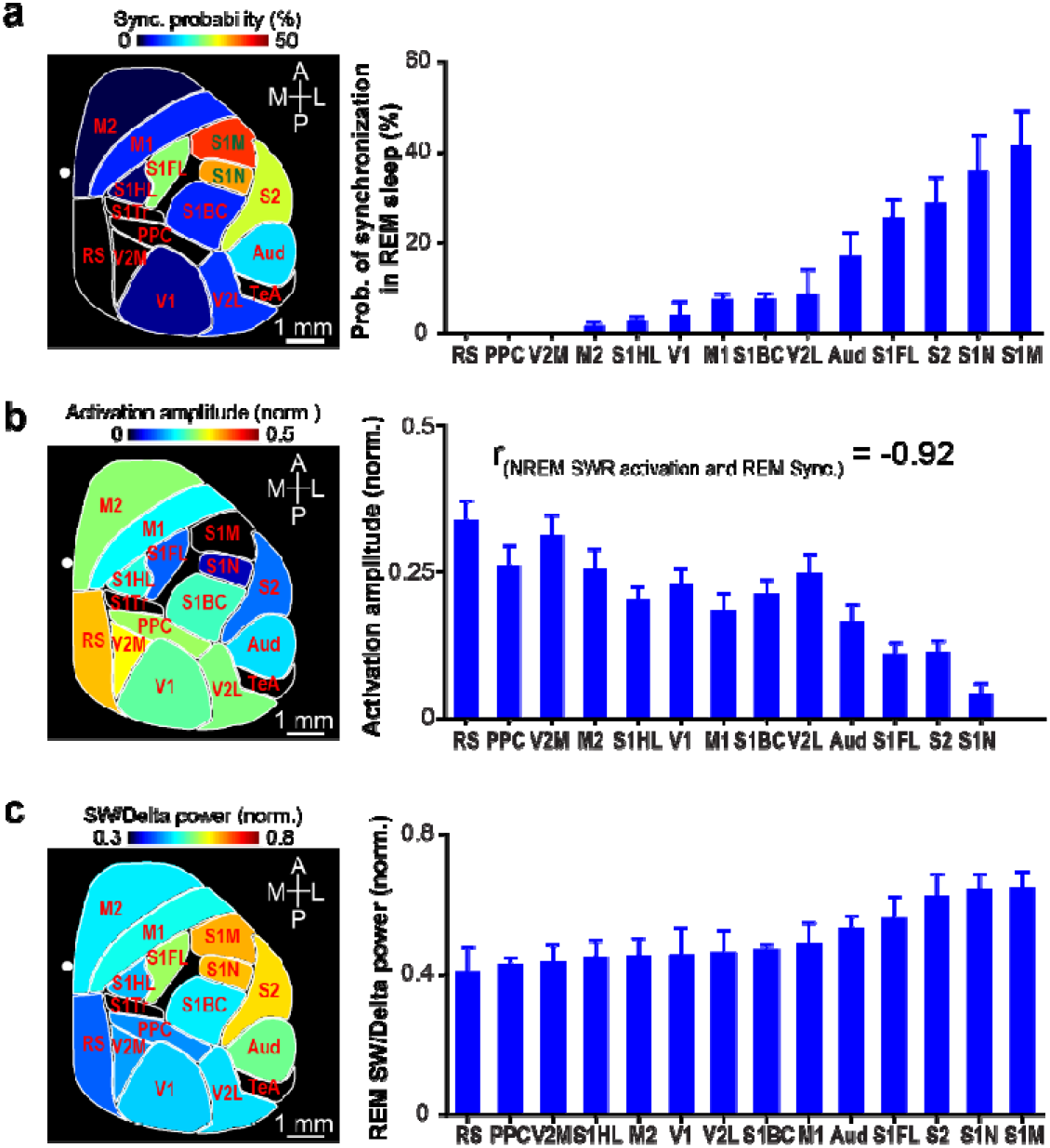
(a) Left: Animal-wise average (n=6 glutamate mice) map shows the probability of synchronized activity during REM sleep across different cortical regions. Right: Ordered probability of synchronization in 14 ROIs. Error bars, s.e.m. (b) Left: Animal-wise average (n=12 mice) map demonstrates cortical activation around hippocampal sharp-wave ripples during NREM sleep. Right: Cortical activation across 13 ROIs sorted based on probability of synchronization in REM sleep. The amplitude of cortical activation in NREM sleep was negatively correlated with the synchronization probability in REM sleep (r=−0.92, p<0.001). Data was obtained from Karimi Abadchi et al, 2020. (c) Left: 2D map shows the distribution of SW/delta power across regions measured during REM sleep. Right: The 14 ROIs were sorted based on the REM sleep SW/delta power. Error bars, s.e.m.

**Supplementary Figure 4.**
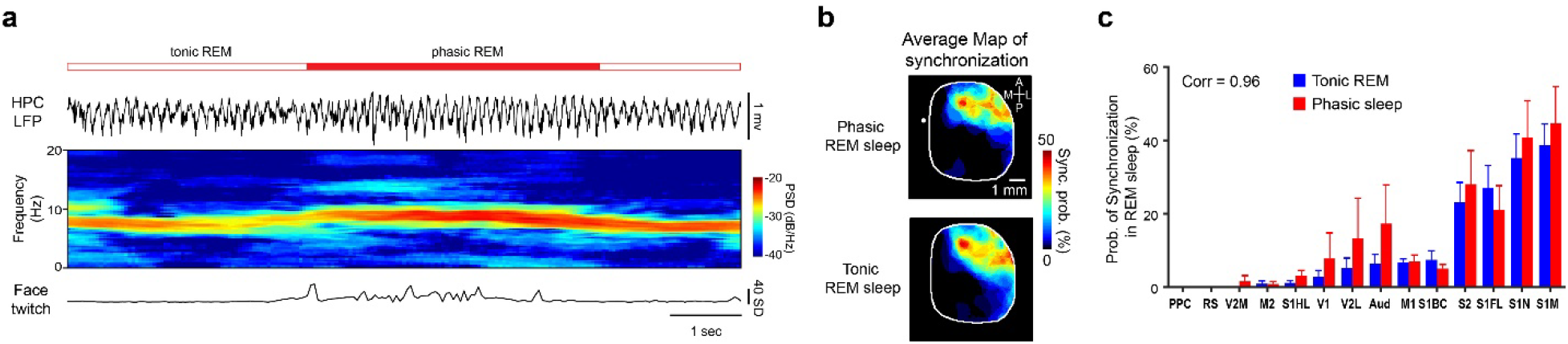
Cortex shows local synchronized patterns during both phasic and tonic REM sleep. (a) Top: Example trace of hippocampal LFP signal showing epochs of phasic and tonic REM sleep in a head-restrained mouse. Middle: Spectrogram of hippocampal LFP illustrates frequency modulation of theta activity across phasic and tonic REM sleep. Bottom: The face movement signal which was used to separate phasic REM events from tonic ones. (b) Epoch-wise average of binarized p-value maps for phasic (top) vs tonic (bottom) REM sleep calculated for a representative animal. Warmer colors show cortical regions with higher probability of synchronization. (c) Bar graph indicates the probability of synchronized activity during phasic and tonic REM sleep across 14 ROIs. ROIs are sorted based on probability of synchronized activity in tonic events. Error bars, s.e.m. (n=5 mice; paired t-test, n.s. p>0.05).

**Supplementary Figure 5.**
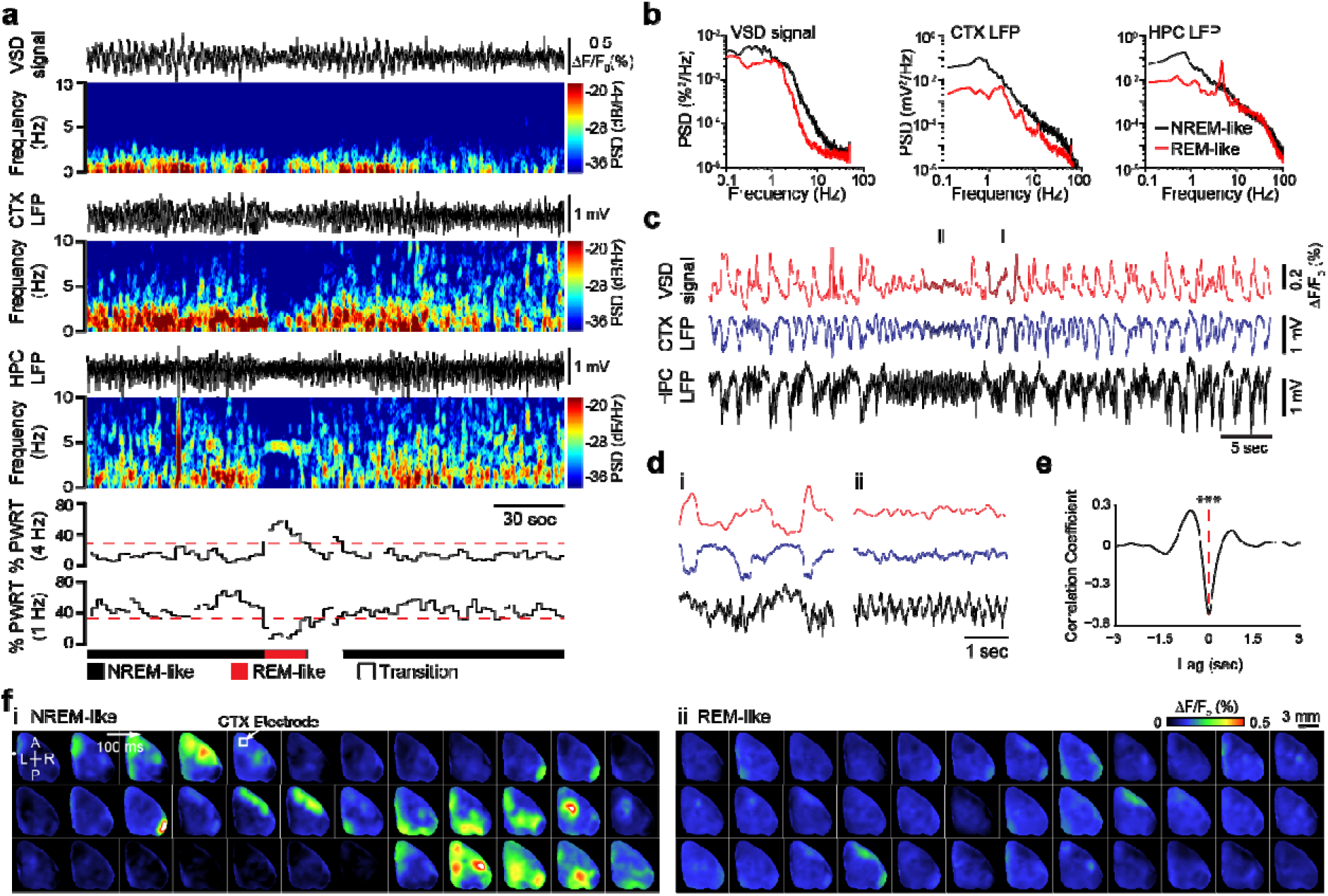
Spontaneous state alternation of brain activity in urethane anesthetized mouse. (a) A cortical VSD (first panel from top), cortical LFP (second panel), and hippocampal LFP traces (third panel), as well as corresponding spectrograms showing spontaneous alternation of brain states during urethane anesthesia. The VSD signal was derived from an ROI (0.112 mm^2^) around the cortical electrode in the primary motor cortex. The percentages of the total power in the 3-5 Hz and 0.2-1.2 Hz bands for hippocampal LFP are displayed in the fourth panel. Their ratio was used to score the recording into NREM-like, REM-like and transition states. (b) Power spectral analysis reveals a reduction of below 1 Hz power in the VSD, cortical (CTX) LFP, and hippocampal (HPC) LFP signals during the REM-like state. (c) Representative traces of VSD (0.2-7 Hz) and cortical and hippocampal LFP signals are shown. (d) Time expansion of signals highlighted in c i and ii are shown. Vertical scale bars are the same as c. (e) Cross-correlogram between 15 minutes of VSD signal and cortical LFP recorded from the same area (M1). Note that two signals are highly correlated. (*** p < 0.001, Student’s t-test). (f) Montage of spontaneous VSD activity (0.2-7 Hz), corresponding to epochs i and ii, demonstrates different spatiotemporal dynamics of cortical activity during NREM-like and REM-like states. Note that diverse cortical regions get activated during NREM-like up states. During REM state, most of the cortex shows desynchronized activity.

**Supplementary Figure 6.**
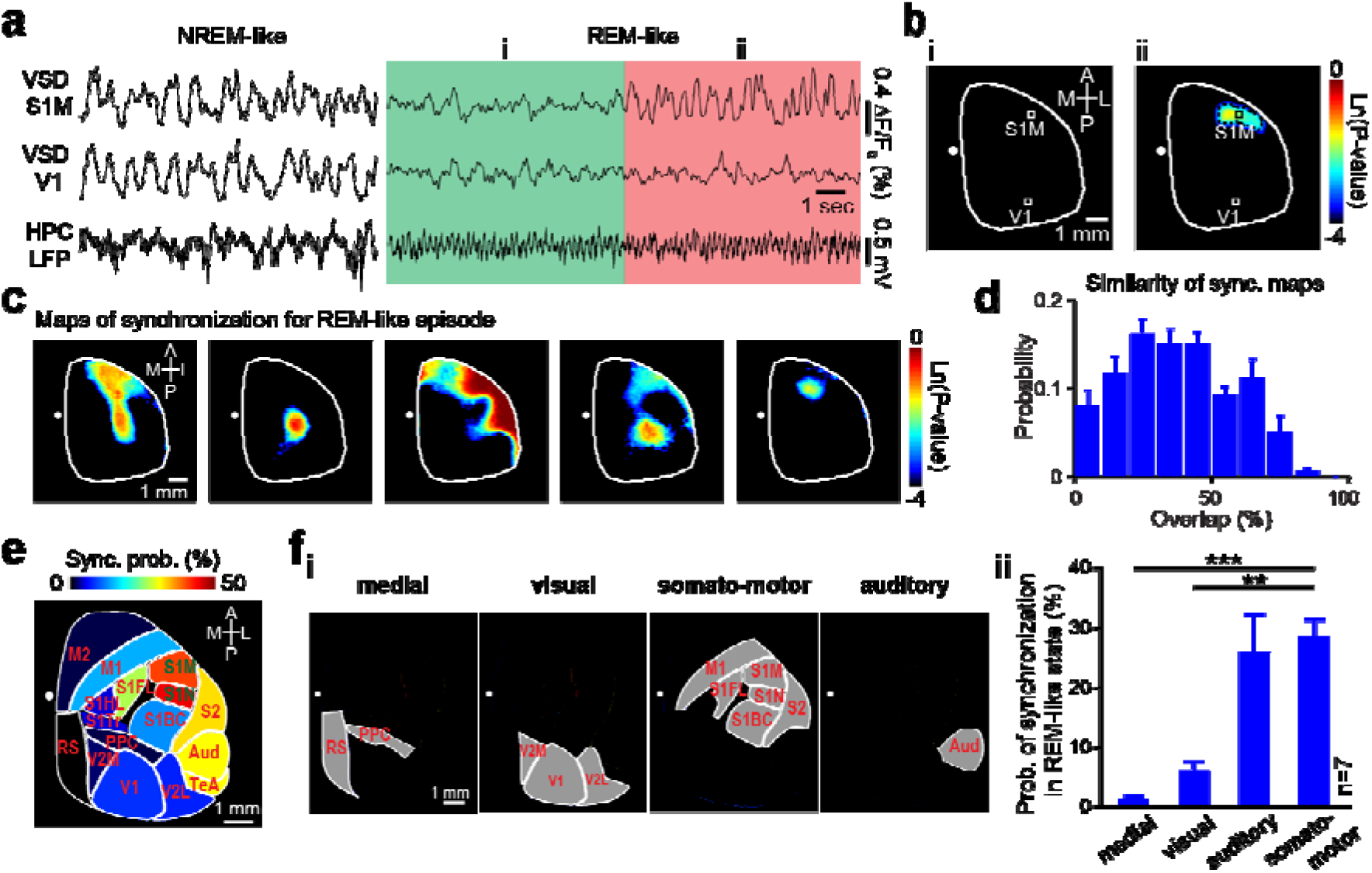
Local cortical synchronization during REM-like state assessed by VSD imaging in urethane anesthetized mice. (a) Traces show VSD signals (top and middle traces) derived from two ROIs (S1M and V1) and simultaneously recorded hippocampal LFP signal during NREM-like (left) and REM-like episodes (right). (b) Representative p-value maps of synchronization calculated for 2 epochs (8 sec long) of a REM sleep episode highlighted in green (i) and red (ii) in a. (c) Similar to b, but for five consecutive 8-sec epochs during a REM-like episode. (d) Histogram of overlap between all pairs of binarized p-value maps pooled across 7 mice. Error bars, s.e.m. (e) Binarized p-value maps from 7 animals were averaged and registered onto the Allen Institute Mouse Brain Coordinate Atlas. (f) (i) Four major structurally defined neocortical subnetworks. (ii) Bar graph shows the probability of synchronized activity during REM-like state across cortical subnetworks, sorted in ascending order. Error bars, s.e.m. (repeated measure ANOVA with Greenhouse-Geisser correction for sphericity: F3,18 = 15.996, p = 0.0021; post-hoc multiple comparison with Tuckey’s correction: medial vs visual p = 0.0590, medial vs auditory p = 0.0379, medial vs somatomotor p = 3.779×10^−4^, visual vs auditory p = 0.0421, visual vs somatomotor p = 2.425×10^−3^, auditory vs somatomotor p = 0.9786).

**Supplementary Figure 7.**
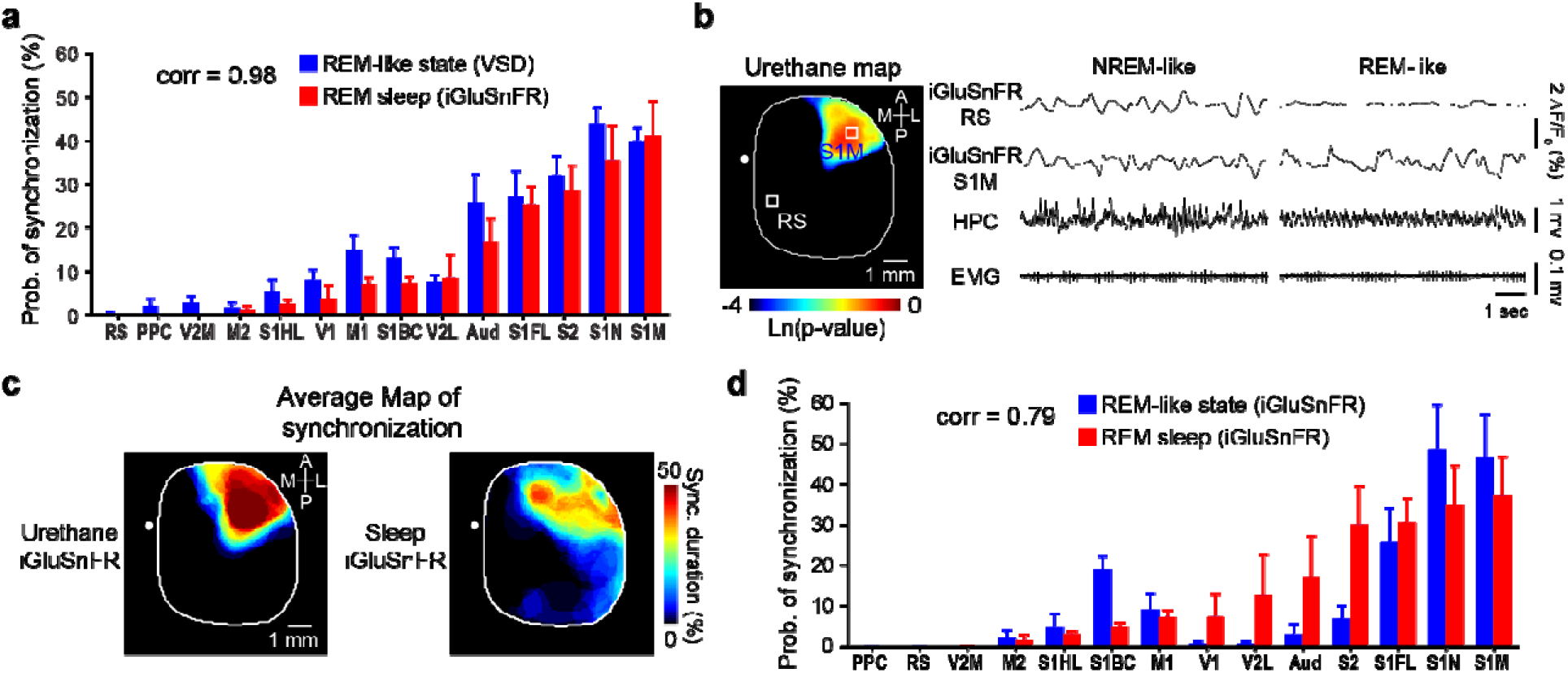
(a) Region-wise comparison of synchronization probability during REM-like state of VSD experiments and REM sleep of glutamate experiments (n=7 and 6 respectively; p-value > 0.05 for all ROIs, student’s t-test). ROIs are ordered based on the synchronization probability in REM sleep. Moreover, the rank order of synchronization probability in REM-like state was similar to the rank order of synchronization probability in REM sleep. Error bars, s.e.m. (p < 0.001, Spearman’s rank correlation). (b) Left, p-value map of synchronization for a representative REM-like epoch recorded using glutamate imaging. Right, cortical activity in retrosplenial and mouth primary somatosensory cortices along with hippocampal LFP and neck EMG recorded during the NREM-like and REM-like states corresponding to the lef*t* p-value map. (c) Maps of synchronization in the REM-like state and REM sleep for a glutamate mouse were juxtaposed for comparison purpose. (d) Comparison of synchronization probability during the REM-like state and REM sleep in glutamate mice (n=3 for both groups). ROIs are ordered based on the synchronization probability in REM sleep. Error bars, s.e.m.

**Supplementary Figure 8.**
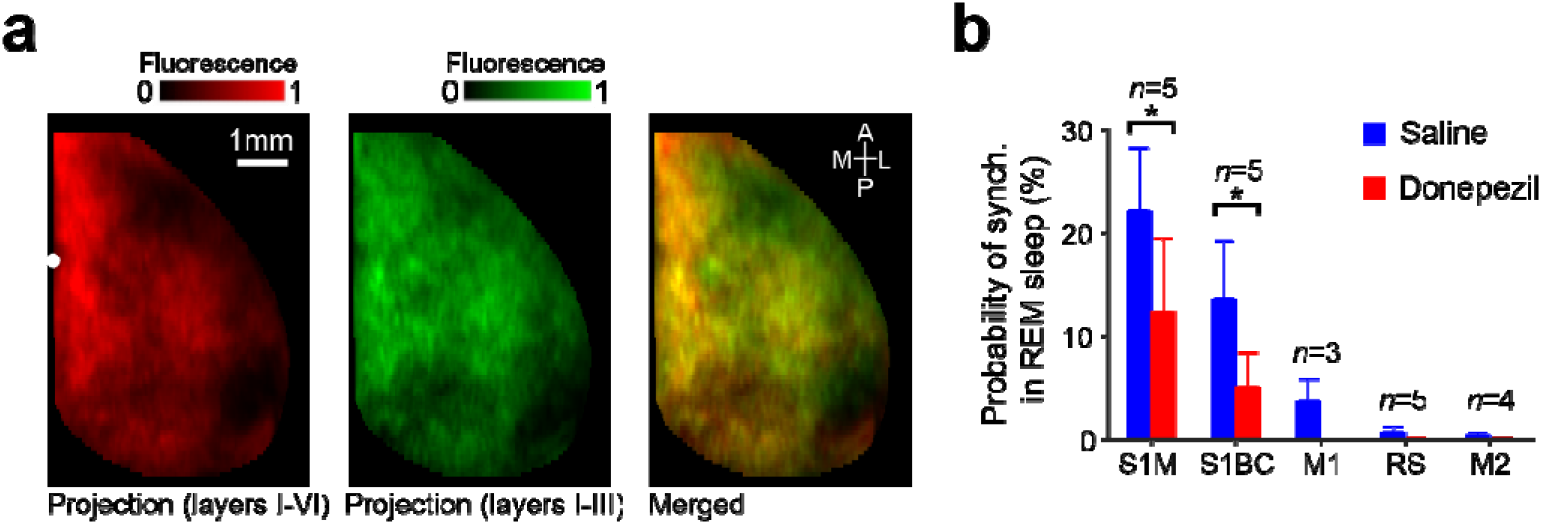
(a) Images illustrate maps of the axonal projections from the basal forebrain to the entire cortical laminae (left), supragranular layers (middle), and their overlay (right) (b) Increase of extracellular acetylcholine level by donepezil (1 mg/kg) significantly reduced slow-wave activity during REM sleep, particularly in the mouth and barrel primary somatosensory regions. Error bars, s.e.m. (paired t-test, *p<0.05).

**Supplementary Figure 9.**
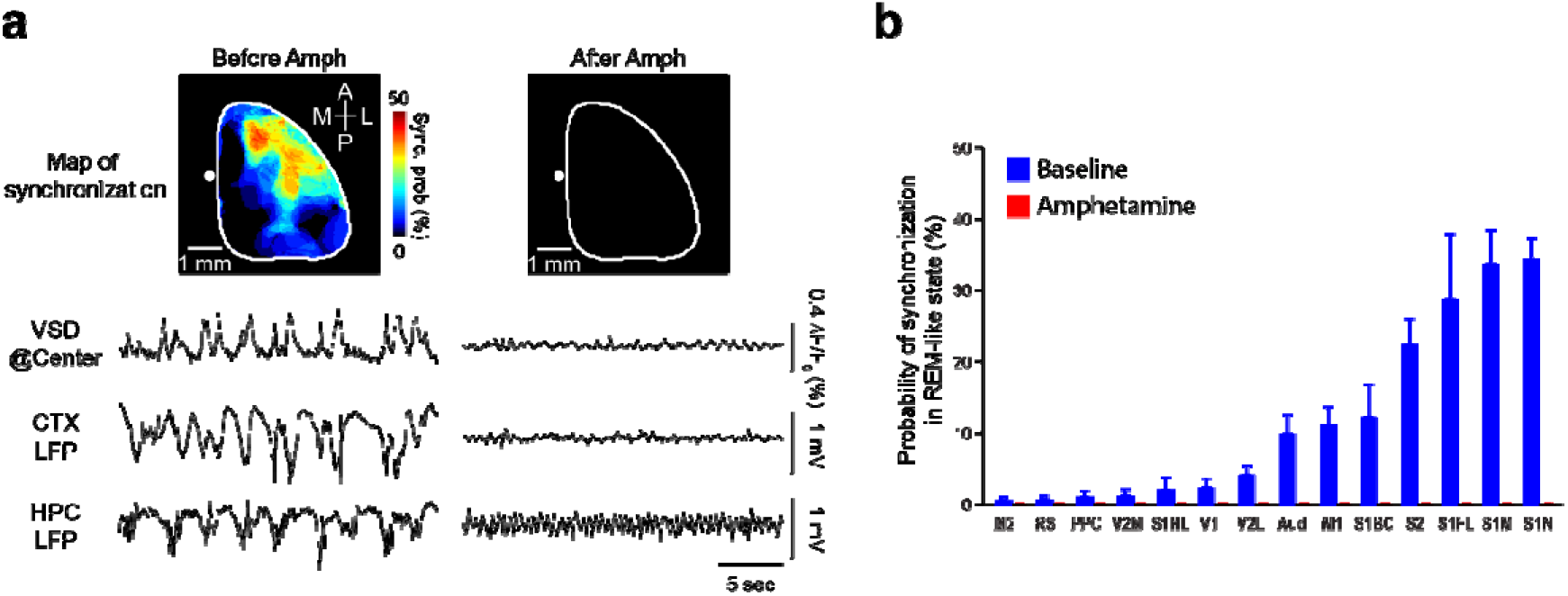
Amphetamine abolishes local SWA during the REM-like state. (a) Top: Effect of amphetamine administration (0.1 mg/kg) on the map of synchronization. Bottom: Representative traces of VSD and cortical and hippocampal LFP signals are shown before and 30 minutes after amphetamine administration. The VSD signal was derived from an ROI (0.112 mm^2^) around the cortical electrode in the primary motor cortex. (b) Comparison of the cortical synchronization probability between the spontaneous REM-like state and the amphetamine-induced neural desynchronization. Error bars, s.e.m. (n=3 mice).

**Supplementary Figure 10.**
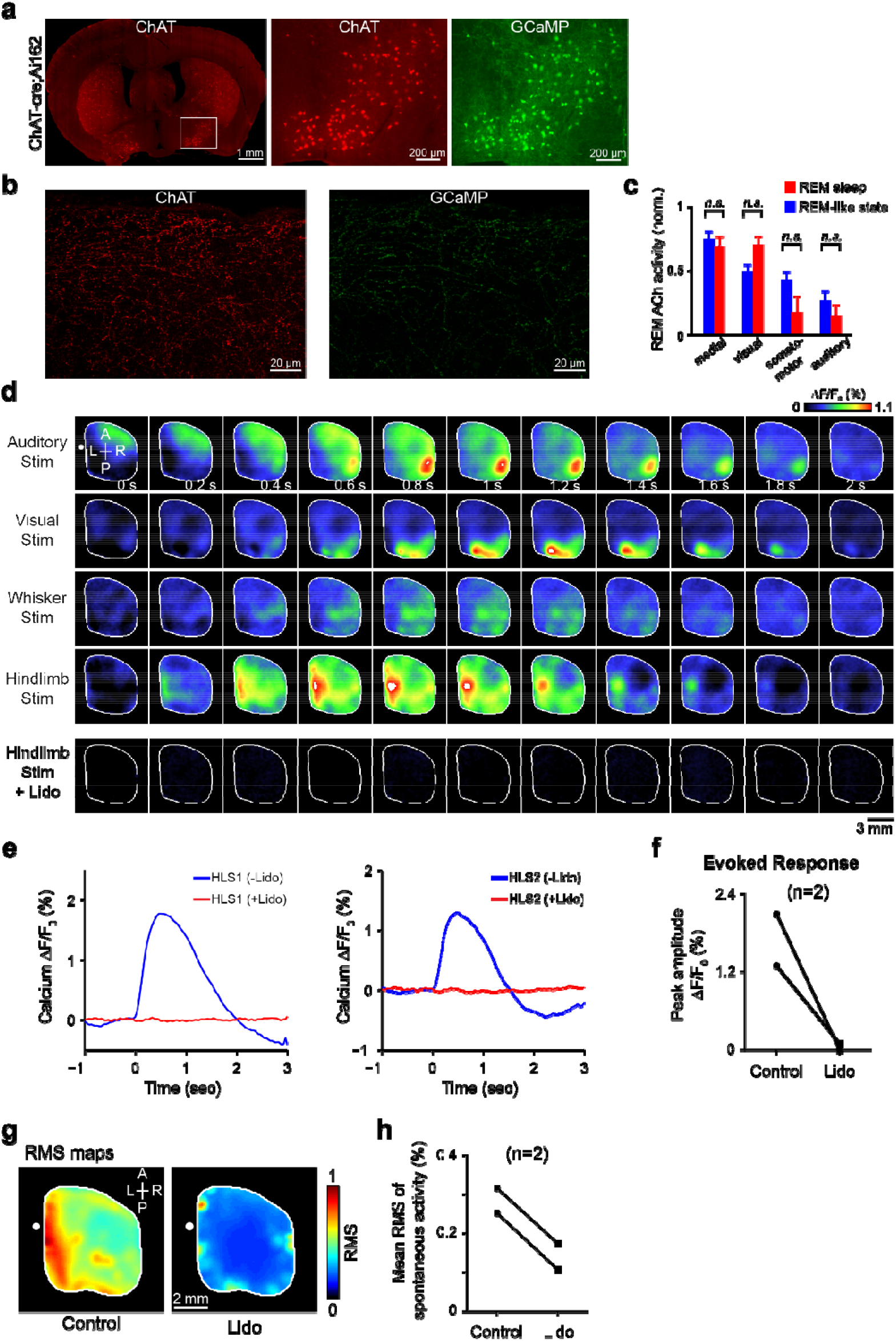
(a) Coronal section of brain showing expression of GCaMP6s in cholinergic neurons of the basal forebrain in ChAT□Cre;Ai162 mice. Red shows anti□choline acetyltransferase (anti-ChAT) immunofluorescence and green shows GCaMP6s. (b) Confocal image of cholinergic axons in the primary visual cortex. ChAT□expressing axons co□labeled with GCaMP6s (enhanced with anti_JGFP staining) in Chat□Cre;Ai162 mice. (c) ACh activity in four cortical subnetworks measured in the urethane-induced REM-like state and REM sleep. Error bars, s.e.m. (n=6 and 4 for urethane and natural sleep respectively, Wilcoxon rank-sum test, *n.s.* p>0.05). (d) Montages of evoked calcium activity following multiple modalities of sensory stimulation, including auditory tone stimulation, visual stimulation, whisker stimulation and hindlimb electrical stimulation. Hindlimb stimulation was repeated after topical application of lidocaine (2%; last row). (e) Time course of hindlimb stimulation evoked activity from 0.112 mm^2^ ROIs chosen from primary (left) and secondary (right) hindlimb somatosensory cortices before (blue) and 10 minutes after (red) lidocaine administration. (f) Comparison of the peak amplitude of evoked calcium activity in HLS1 between the control and lidocaine groups. (g) Root mean square (RMS) maps of spontaneous activity (~ 20 min) prior to and following the application of lidocaine (2%). (h) Pixel-wise average of RMS maps for two animals before and after lidocaine administration.

## Notes

### Competing Interest Statement

The authors have declared no competing interest.

